# Engineered minimal type I CRISPR-Cas system for transcriptional activation and base editing in human cells

**DOI:** 10.1101/2024.01.26.577312

**Authors:** Jing Guo, Luyao Gong, Haiying Yu, Ming Li, Zhenquan Liu, Shuru Fan, Changjialian Yang, Dahe Zhao, Jing Han, Hua Xiang

## Abstract

Type I CRISPR-Cas systems are widespread and have exhibited remarkable versatility and efficiency in genome editing and gene regulation in prokaryotes. However, due to the multi-subunit composition and large size, their application in eukaryotes has not been thoroughly investigated. Here, we demonstrate that the type I-F2 Cascade, the most compact among type I systems and significantly smaller than SpCas9, can be developed into programmable tools for use in human cells. For transcriptional activation, the efficiency of the tool based on the engineered I-F2 system can match or surpass that of dCas9. Besides, narrow editing windows limit the application of base editors. Although the R-loop formed by Cascade is much wider than that by Cas9 or Cas12, the potential of base editing with Cascade has not yet been explored. We successfully created a base editor with the I-F2 Cascade, which induces a considerably wide editing window (∼30 nt) with a bimodal distribution. The wide editing window can expand the range of targetable sites and can be useful for disrupting functional sequences and genetic screening. The editing efficiency can achieve 50% in human cells. This research underscores the application potential of compact type I systems in eukaryotes and developed a new base editor with an extraordinary wide editing window.

## Introduction

CRISPR-Cas systems are adaptive immune systems that are widely distributed in prokaryotes. They can be classified into two classes, six types, and more than 30 subtypes^1, 2^. Class 1 encompasses types I, III, and IV, with effector modules composed of multiple proteins. Class 2 comprises types II, V, and VI, characterized by a single, multi-domain effector protein. The CRISPR immunity mechanism typically involves three processes: adaptation, crRNA biogenesis, and interference^3^.

The type I system is the most prevalent type of CRISPR-Cas and is divided into seven subtypes, I-A to I-G^1^. The crRNA-binding complex of type I is called Cascade (CRISPR-associated complex for antiviral defense), which binds to the crRNA for target recognition, facilitates duplex formation between the crRNA and its complementary target DNA for R-loop formation, and recruits Cas3 nuclease to degrade DNA processively^4–7^. Type I systems have been developed into various microbial genome manipulation tools, such as tools for genome editing with homologous repair templates^8–10^, transcriptional regulation^11, 12^, large fragment deletion^13, 14^, large fragment integration without requiring homologous recombination^15^, and simultaneous genome editing and gene regulation with Cascade-Cas3^16^. Since 2019, several type I subtypes have been successfully used in eukaryotic genome manipulation. When Cascade and Cas3 were expressed simultaneously, Cas3 exhibited unique exonuclease activity, resulting in large fragment deletions of up to 200 kb^17–23^. Similar activity was reported for type I-D, of which Cas10 is a functional nuclease, while short indels were detected as well^24, 25^. Fusing Cas3 with the cytidine deaminase enables wide-ranging random mutagenesis, covering up to 55 kb, which provides an efficient tool for optimizing complex biosynthetic pathways^26^. Moreover, Cascade can be fused with different domains to perform different functions. When Cascade was fused to the dimerization-dependent, non-specific FokI nuclease domain, the editing efficiency was comparable to that of dCas9-FokI^17^. When Cascade was fused to the transcriptional activation or repression domain, the expression levels of targeted endogenous genes could be modulated^27–29^.

However, the Cascades used in eukaryotes comprise four or five subunits, counting in Cas6, which processes the precursor crRNA (pre-crRNA) into mature crRNA. For example, the Cascade of *Escherichia coli* (*E. coli*) K12 type I-E system is composed of five subunits, with a total gene size of ∼4.4 kb^19, 29^. The Cascade of *Neisseria lactamica* ATCC 23970 type I-C system is composed of four subunits, with a total gene size of ∼3.8 kb, including the hidden small subunit ^21^. The large size hinders applications with cargo size constraints, such as the widely applied adeno-associated virus (AAV) vector with a ∼4.7 kb packing limit, which hampers most CRISPR tools’ clinical applications^30–33^. Thus, mining more compact type I systems will be helpful for applying type I genome manipulation tools to eukaryotes.

On the other hand, R-loop formation is an important feature of CRISPR systems. In the R-loop structure, the non-target DNA strand displaced by the spacer of crRNA is exposed and accessible to other molecules, which is exploited by base editing technologies^34–36^. By fusing single-stranded DNA (ssDNA) deaminases or glycosylases to catalytically impaired Cas nuclease, base editors can install various base conversions without the need for double-stranded DNA (dsDNA) breaks or donor DNA templates^36,37^. However, base editors are developed mainly with catalytically impaired Cas9 or Cas12 (including their presumed compact ancestors)^36, 38–41^. It is noteworthy that the R-loop formed by type I systems is considerably wider, spanning around 30–40 nt^5, 20, 34^, compared to that in Cas9 or Cas12, which has a narrower R-loop of approximately 20 nt^34, 35, 42–44^. This extended R-loop in type I systems may provide more nucleotide substrates accessible to ssDNA deaminases or glycosylases, thus expanding targetable sites. Therefore, exploring the potential of type I systems in base editing could be valuable for developing new base editors with unique editing windows.

Type I-F2 systems feature the most compact Cascade identified to date, consisting of Cas5, Cas6, and Cas7 without the large subunit (Cas8) or small subunit (Cas11)^45–47^. The total gene size is significantly smaller than that of SpCas9^42, 48–51^, making it a promising candidate for easier delivery in eukaryotes. However, previous attempts to apply this system to eukaryotes were unsuccessful. For instance, the Cas3 and Cas5-7 derived from the *Shewanella putrefaciens* CN32 (Spu) I-F2 did not exhibit DNA cleavage activity in human cells^19^. Similarly, another study reported that the Cas5-7 derived from Spu I-F2 could not activate the expression of the target gene, after fusing to a transcriptional activator^27^. Here, we present a compact type I-F2 system for manipulating the human genome. The Cascade is derived from the *Moraxella osloensis* CCUG 350 (Mos350), with a total gene size of approximately 2.7 kb, and prefers a simple 5′-CC protospacer adjacent motif (PAM). By fusing the transcriptional activation domain with the Cascade, we can modulate gene expression in human cells and achieve robust activity through crRNA and Cascade engineering. Additionally, by fusing deoxyadenosine deaminase with the Cascade, we developed a base editor with the type I-F2 system. Interestingly, this adenine base editor (ABE) induces a uniquely wide editing window (∼30 nt) with a bimodal distribution and can achieve an editing efficiency of up to 50%. The I-F2 ABE with a wide editing window can be useful for genetic screening and disrupting the functional sequences. These results highlight the potential of compact type I-F2 systems in eukaryotes and expand the base editing toolbox.

### Identification of miniaturized type I-F2 CRISPR-Cas systems

Type I-F2 systems consist of five Cas proteins: Cas1, Cas3, Cas5, Cas6, and Cas7, without the large and small subunits (Fig. 1a)^45–47^. Cas1, Cas3, and Cas6 of type I-F2 have sequence similarities to those of type I-F1 or I-F3 systems, while Cas5 and Cas7 of type I-F2 have no significant sequence homology to any other type I subtypes^46, 47^. Previously, studies of type I-F2 focused on the Spu I-F2 system, while this system is proven disabled in gene activation^27^ or genome editing in human cells^19^. To develop genetic tools with compact type I systems, we searched public databases for miniaturized type I-F2 systems and found out 93 systems (Supplementary Fig. 1 and Supplementary Table 1). Phylogenetic analysis of Cas7 showed that Cas7 of types I-F1 and I-F3 clustered together, whereas all 93 systems of type I-F2 clustered in a separate branch (Fig. 1b), confirming strong sequence deviations in the Cas7 homologs. The gene sizes of these I-F2 Cascades are approximately 2.4–2.8 kb. The repeats are 28 bp long, and the sequences are highly conserved according to multiple sequence alignments (Supplementary Fig. 2a). The spacer of 32 bp accounted for 96.6% of all the spacers (Supplementary Fig. 2b and Supplementary Table 2), and the wild-type spacers used in subsequent experiments are of 32 bp. Interestingly, the average number of spacers is 55, ranging from 3 to 180, indicating that I-F2 systems are highly active.

**Fig. 1:**
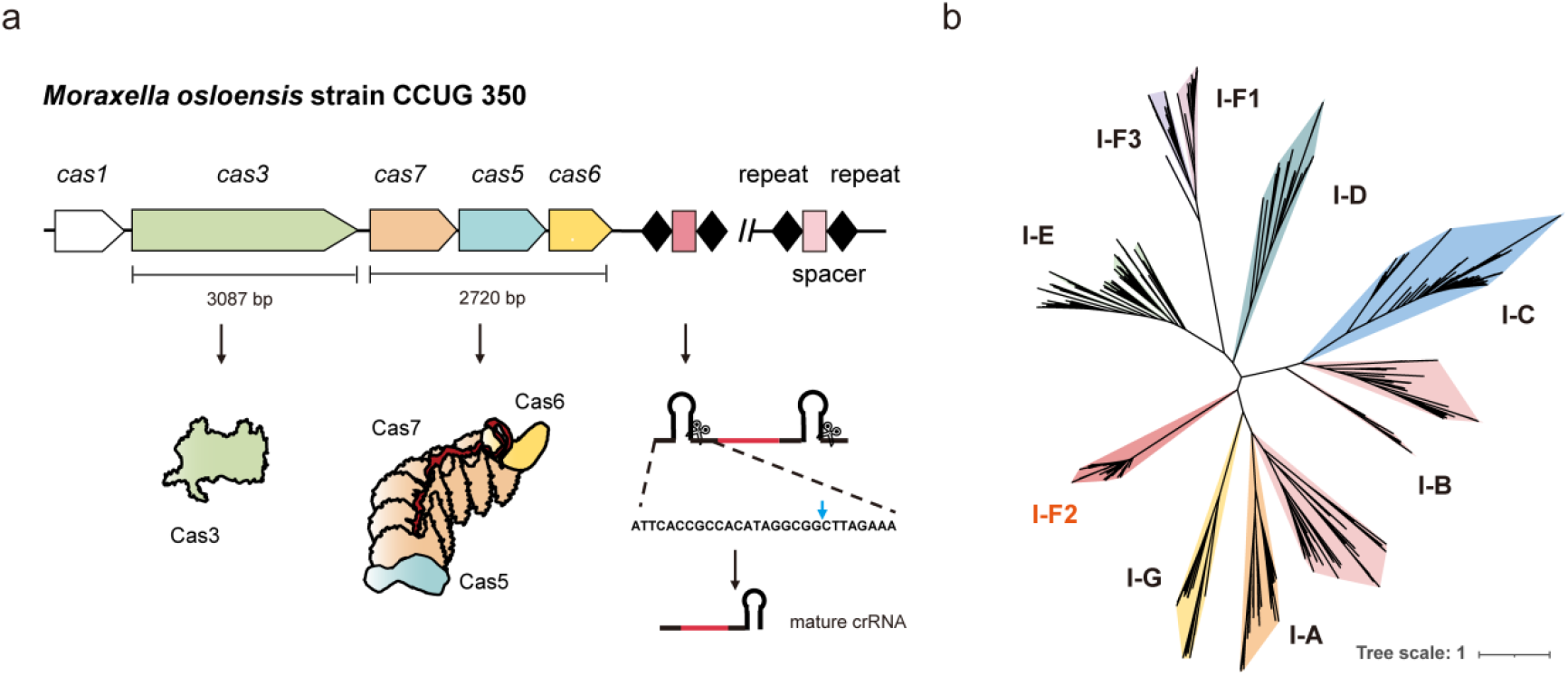
The minimal type I-F2 CRISPR-Cas system. **a,** Schematic representation of the *Moraxella osloensis* strain CCUG 350 type I-F2 CRISPR-Cas system showing Mos350 Cascade stoichiometry and crRNA processing. The genes comprising the Mos350 Cascade complex are represented in different colors. Scale bars show the gene sizes of Cas3 and Cascade (including intergenic regions). Repeats and spacers in the CRISPR array are indicated using black diamonds and red rectangles, respectively. Cas6 cleaves pre-crRNA transcripts at the positions indicated by the blue arrow in the repeat sequences to produce mature crRNAs. **b,** The identified I-F2 Cas7 proteins were subjected to phylogenetic analysis using the maximum likelihood method, along with Cas7 proteins from other type I systems. The scale bar represents the number of substitutions per site.

### Targeted gene repression by I-F2 Cascades in *E. coli*

First, we identified the characteristics of type I-F2 systems in *E. coli*. To analyze the PAM preference, we conducted a PAM depletion assay and plasmid interference assay using the Mos350 I-F2 system. A plasmid library with four randomized nucleotides at the 5′ end of the protospacer was constructed. The results reveal that this system is capable of crRNA-guided dsDNA targeting in *E. coli* and prefers a simple 5′-CC PAM (Fig. 2a and Supplementary Fig. 3).

**Fig. 2:**
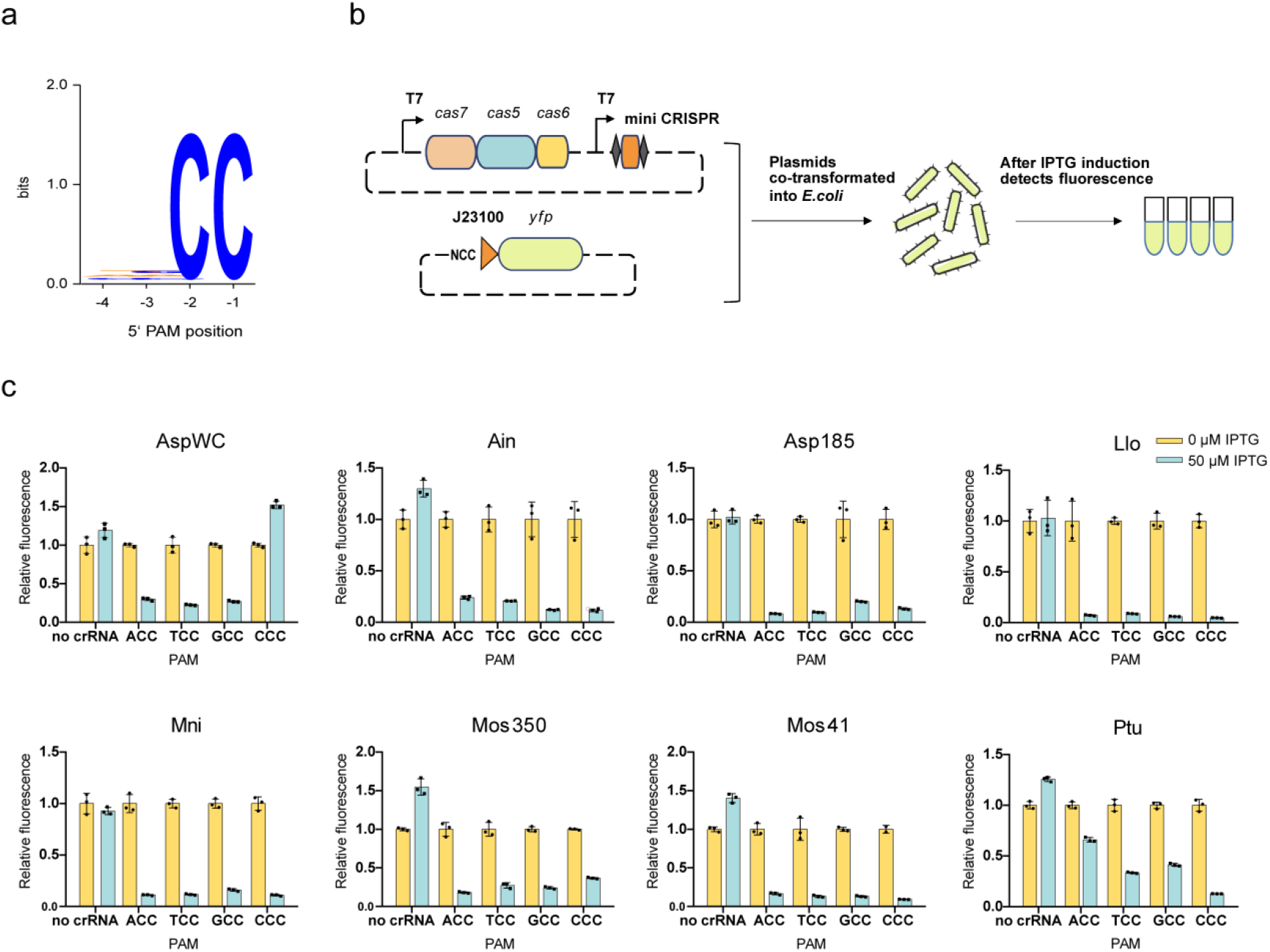
Targeted gene repression by I-F2 Cascades from eight different microbes in *E. coli*. **a,** WebLogos for the PAM of the Mos350 Cascade determined using plasmid depletion assays. **b,** CRISPR-mediated gene repression scheme. Cascade-crRNA targeted the YFP promoter with the NCC PAM motif at the 5′ end. **c,** Changes in the relative YFP fluorescence with the expression of I-F2 Cascades from eight different microbes (Supplementary Fig. 4). No crRNA: The plasmid expressing the Cascade (without the mini-CRISPR) and the target plasmid were co-transformed. All values are represented as mean ± s.d. of three biological replicates.

Since the Cascade-crRNA complex is responsible for target recognition and R-loop formation, it can be a useful component for gene regulation or base editing tools. We tested the targetable DNA-binding ability of type I-F2 Cascades in *E. coli*. Except for Mos350, we synthesized seven additional type I-F2 systems from different strains (Supplementary Fig. 4). We designed the NCC PAM at the 5′ end of the protospacer (in the promoter region of YFP) (Fig. 2b). After inducing the expression of Cascade and crRNA, significant decreases in fluorescence intensity were observed for all eight I-F2 systems, by recognizing 5′-CC PAM, except for the system derived from *Acinetobacter* sp. WCHAc010052 (AspWC), which prefers 5′-DCC PAM (Fig. 2c). These results indicate that all eight I-F2 Cascades have programmable DNA-binding abilities in *E. coli*.

### Expression and complex formation of I-F2 Cascades in human cells

To repurpose miniaturized I-F2 Cascades for application in human cells (HEK293T cells), we used the cytomegalovirus (CMV) promoter to express the three subunits of Cascade (human codon-optimized Cas5/Cas6/Cas7), which were linked by self-cleaving 2A peptides. Nuclear localization signals (NLS) derived from the simian virus 40 (SV40) large T antigen were fused to each N-terminus of the three subunits. The RNA polymerase III U6 promoter was used to transcribe the pre-crRNA, which could be processed by Cas6 into a mature crRNA (Fig. 3a).

**Fig. 3:**
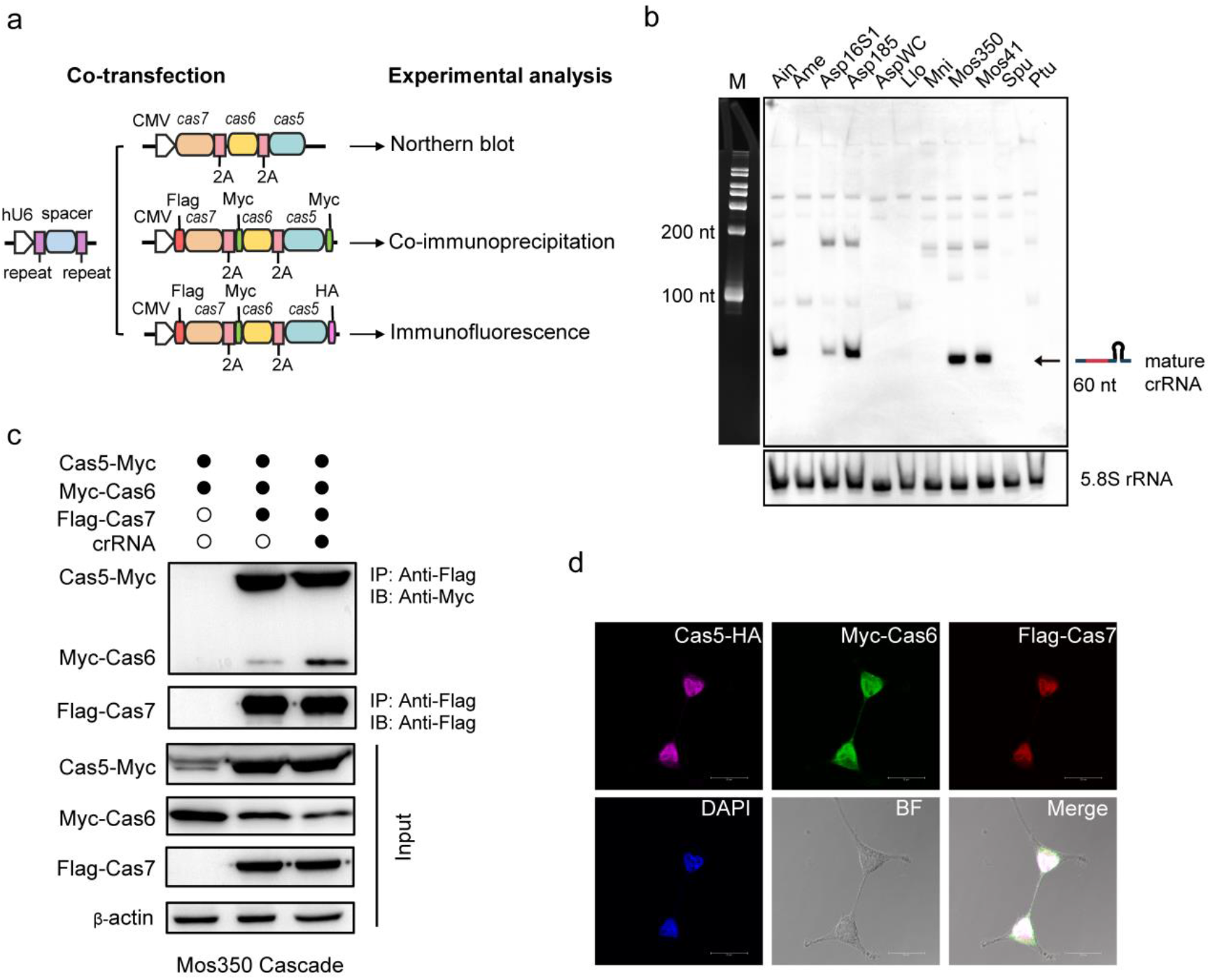
Characterization of the crRNA processing and Cascade complex formation of type I-F2 in human cells. **a,** The experimental design is illustrated as follows: the U6 promoter on the plasmid drives crRNA expression and the CMV promoter drives Cas5, Cas6, and Cas7 expression. The two plasmids were co-transfected into HEK293T cells. **b,** Northern blot assays to demonstrate the processing of crRNAs by I-F2 Cascades from 11 different strains (Supplementary Fig. 4) with the same spacer-specific probe and 5.8S rRNA was probed as the inner control in HEK293T cells. **c,** Co-immunoprecipitation and Western blot showed the formation of the Cascade complex. The proteins were tagged as shown in (a). IB, immunoblot; IP, immunoprecipitation. **d,** Immunofluorescence assay to detect nuclear localization signals. Red, green, and rose illustrate the Cas7, Cas6, and Cas5 subunits, respectively. The nucleus was stained using DAPI. BF showed the cellular morphology. The previous five photos were merged to yield the results of the nucleus localization of the Mos350 Cascade. For (**b-d),** two independent experiments indicated similar results.

We synthesized 11 type I-F2 systems derived from different strains (Supplementary Fig. 4). Northern blot assays were performed to determine whether these crRNAs could be processed and stabilized in human cells. The result showed that only five systems generated mature crRNAs (Fig. 3b). The mature crRNAs of the Spu I-F2 system were not detected. These five systems were used for further studies. Then, we performed small RNA sequencing to determine the mature crRNA transcripts of the Mos350 I-F2 system. The result showed that the pre-crRNA was processed into mature crRNAs of 60 nt, which were composed of a full-length spacer flanked by a repeat-derived 5′ handle of 8 nt and a repeat-derived 3′ handle of 20 nt (Supplementary Fig. 5), in consistent with the result of the Spu I-F2 crRNA extracted from Spu strain^46^.

Next, we examined the expression and complex formation of type I-F2 Cascades in human cells using the Mos350 system. The plasmid expressing the Cascade components and the plasmid expressing crRNAs were co-transfected into HEK293T cells. Western blot results showed that all three Cas proteins were expressed (Fig. 3c). Co-immunoprecipitation by pull-down of the Flag epitope on Cas7 demonstrated the formation of the Cascade complex. Moreover, in the absence of the crRNA, the binding affinity between Cas6 and Cas7 was significantly reduced, suggesting that Cas6 could bind to Cascade through interaction with the crRNA (Fig. 3c). In contrast, the binding affinity between Cas5 and Cas7 did not change regardless of the presence or absence of the crRNA. In addition, immunofluorescence analysis confirmed that each of the three subunits could enter the nucleus via the SV40 NLS (Fig. 3d).

### Programmable transcriptional activation by Mos350 Cascade-VPR

The I-F2 Cascade consists of three Cas subunits assembled at various stoichiometries (inferred to be Cas5_1_Cas7_6_Cas6_1_) when bound to the wild-type crRNA^45^, providing multiple options for synthetic fusion with regulatory domains (Fig. 4a). Next, we repurposed the compact I-F2 Cascade for the transcriptional activation of endogenous genes in human cells by fusing with the VP64-p65-Rta (VPR) transcriptional activation domains^27, 52^.

**Fig. 4:**
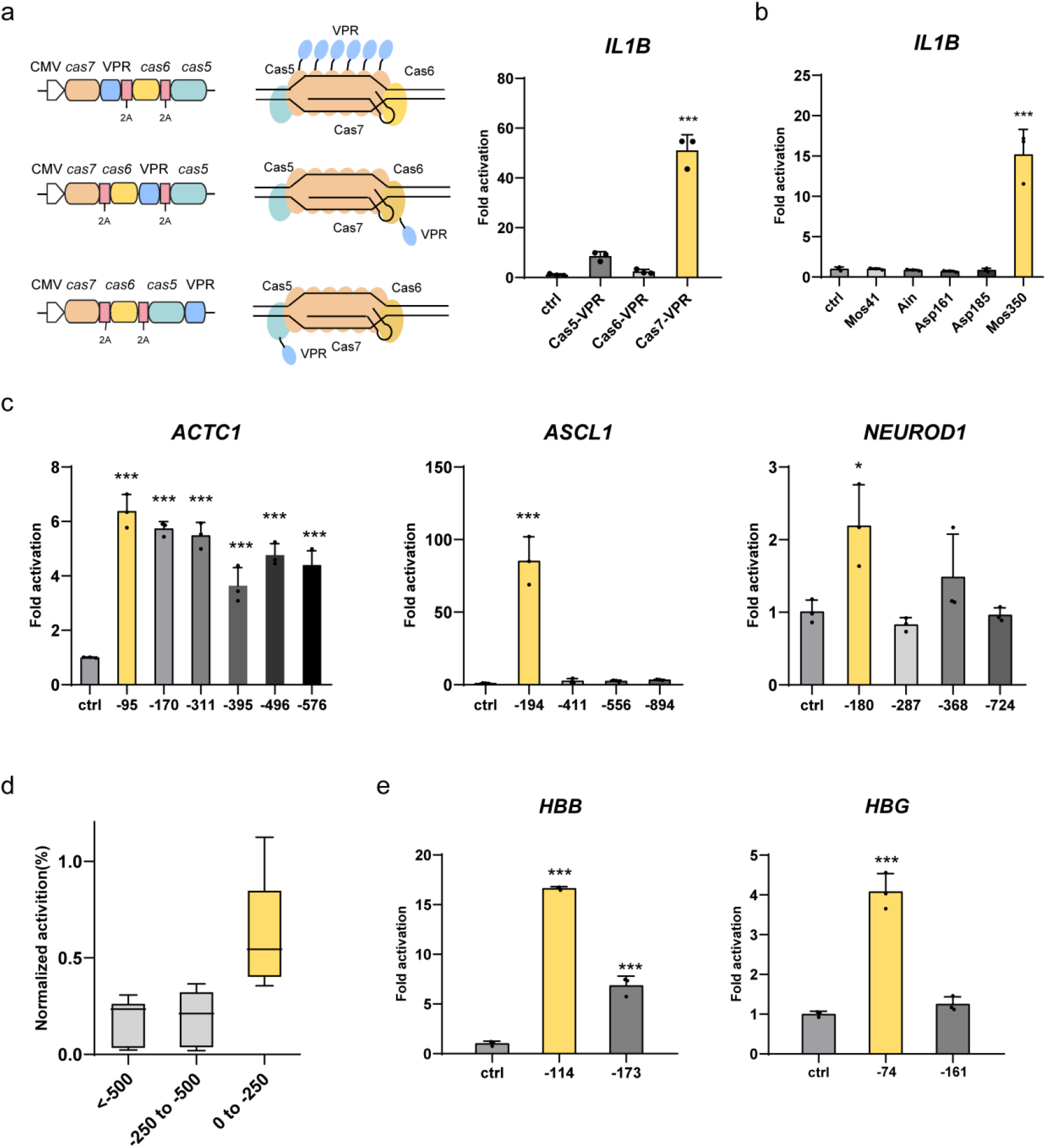
Programmable transcription activation by Mos350 Cascade-VPR. **a,** Left: Gene architectures of VPR transcriptional activators fused with different subunits of the I-F2 Mos350 Cascade. Middle: I-F2 Mos350 Cascade with different subunits fused to VPR transcriptional activators may result in different locations and copy numbers of VPRs. Orange: Cas7; Yellow: Cas6; Cyan: Cas5; Blue: VPR. Right: Fold activations of the *IL1B* gene by different VPR fusion approaches. **b,** Fold activations of the *IL1B* gene by Cascades from different strains with Cas7 fused to VPR. **c,** Fold activations of three genes (*ACTC1*, *ASCL1*, and *NEUROD1*) when the I-F2 Mos350 Cascade-VPR (Cas7-VPR) targets different regions upstream of the TSS. **d,** Histograms showing normalized mean transcriptional activation levels of *ACTC1*, *ASCL1*, and *NEUROD1*, when I-F2 Mos350 Cascade-VPR (Cas7-VPR) transcriptional activator targeted different regions upstream of the TSS. Normalized values for all 3 genes are plotted as box-and-whisker plots with min to max as an option. Maximum and minimum values are shown as whiskers. The boxes in the graph display the median value and the lower and upper quartiles. **e,** Fold activations of the *HBB* and *HBG* genes with I-F2 Mos350 Cascade-VPR (Cas7-VPR) targeting 0 to 250 bp upstream of the TSS. In (**a**) and (**b**), ctrl: None plasmid was transformed. In (**c**) and (**e**), ctrl: Only the plasmid expressing the Cascade-VPR was transformed. All values are represented as mean ± s.d. of three biological replicates. Internal reference gene (**a-c**) and (**e**): *GAPDH*. Statistical significance was assessed by one-way ANOVA (ns, not significant; *P < 0.05; **P < 0.01; ***P < 0.001).

To test the activation effects of VPRs tethering to different subunits, we expressed the Mos350 Cascades with VPR fused to the C-terminus of Cas5, Cas6, or Cas7 (Fig. 4a). The plasmid expressing the crRNA targeting the promoter of the *IL1B* gene was co-transfected into HEK293T cells. Quantitative real-time PCR (qPCR) results showed that VPR fused to Cas7 significantly activated the expression of *IL1B* (51.0-fold relative to control cells), which could be attributed to the multi-copy form of Cas7-VPR in Cascade (Fig. 4a). The following experiments were performed using the Cas7-VPR fusion strategy. Next, we investigated the DNA-binding activity of the remaining four I-F2 Cascades, which could produce mature crRNAs in human cells. QPCR revealed that only Mos350 Cascade-VPR activated the expression of *IL1B* significantly (Fig. 4b). Thus, the Mos350 I-F2 system was used in subsequent studies. Furthermore, we tried to optimize the linker between Cas7 and VPR and attempted to fuse VPR to the N-terminus of Cas7. None of these attempts significantly enhanced the transcriptional activity (Supplementary Fig. 6). Therefore, we retained the original design of Mos350 Cascade-VPR in subsequent studies.

To determine the programmable endogenous gene activation of the Cascade-VPR complex at different loci in the human genome, a series of crRNAs were designed to target different promoter regions of three genes (*ACTC1*, *ASCL1*, and *NEUROD1*). Mos350 Cascade-VPR activated the expression of all three endogenous genes to varying degrees (Fig. 4c). The highest activities were mostly achieved when targeting the region within 250 bp upstream of the transcriptional start site (TSS) (Fig. 4d), which is consistent with other Cascade-VPR vehicles^27, 29^. Next, we designed crRNAs targeting the region within 250 bp upstream of the TSS in two other genes (*HBB* and *HBG*) and observed gene activation as well (Fig. 4e).

### Enhancement of transcriptional activation through crRNA and protein engineering

To investigate the potential of the Mos350 Cascade-VPR as a more efficient transcriptional activation tool, we optimized both the crRNA and the protein components. With each 6 nt increment in the spacer length of the crRNA, an additional Cas7 subunit can be incorporated into the Cascade complex^45, 53, 54^ (Fig. 5a), which may increase the copies of VPR and lead to stronger activation. We explored the impact of spacer length on Mos350 Cascade-VPR activity at the *IL1B* and *ASCL1* loci, with spacer lengths varying by multiples of six. As the spacer length increased, the level of transcriptional activation gradually increased at both loci. The maximum transcriptional activation effects were achieved when the spacer length reached 50 nt, resulting in increased activity of up to 5.8-fold and 8.6-fold, compared to that using the wild-type spacer of 32 nt, at the *ASCL1* and *IL1B* loci, respectively (Fig. 5b).

**Fig. 5:**
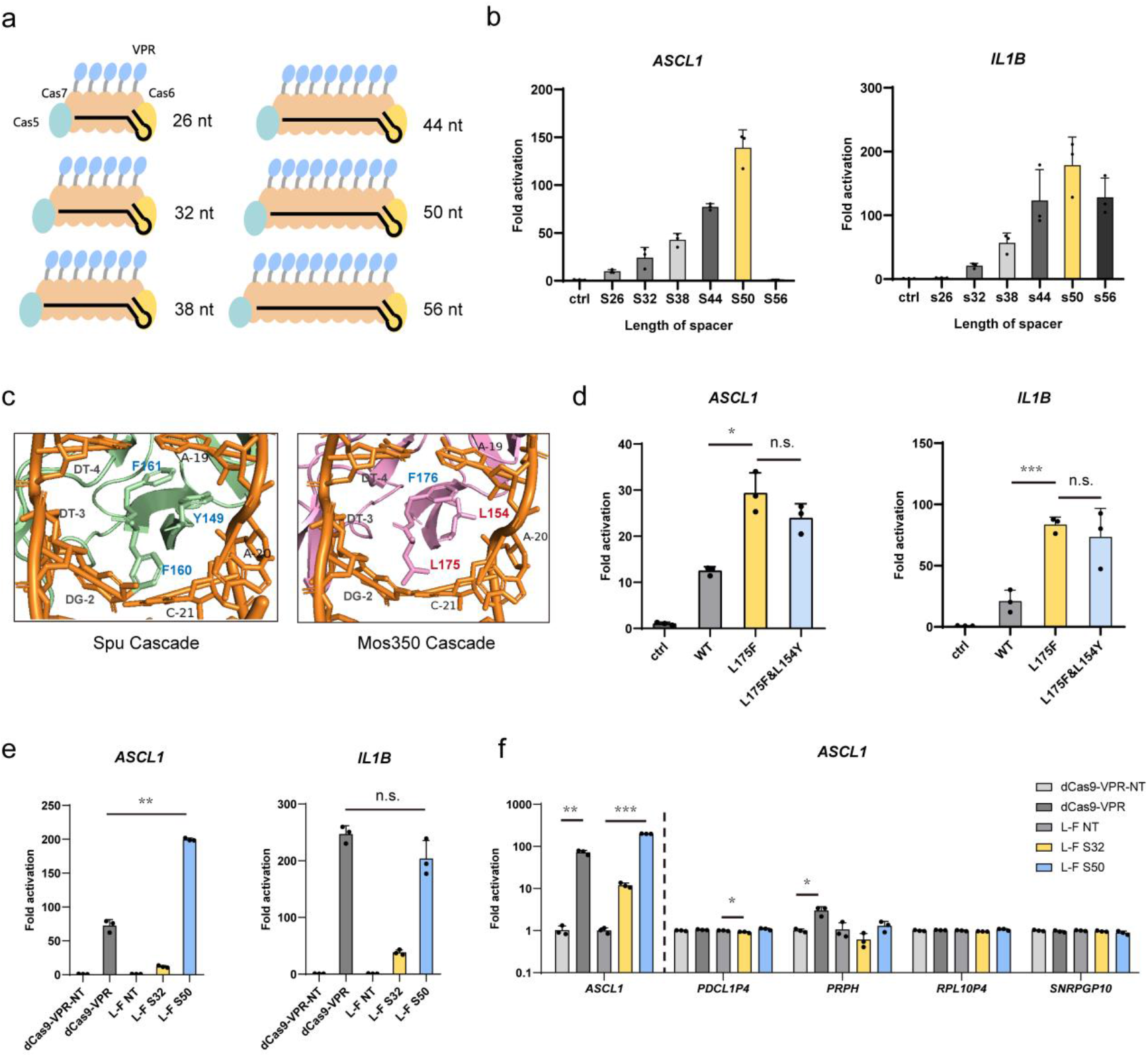
Enhancing transcription activation through crRNA engineering and protein engineering. **a,** Schematic illustration of the speculated structures of Cascade-VPR with various spacer lengths. Orange: Cas7; Yellow: Cas6; Cyan: Cas5; Blue: VPR. **b,** Fold activations of *ASCL1* and *IL1B* genes by Mos350 Cascade-VPR with various spacer lengths. **c,** Left: The crRNA-DNA hybridization double strand was stabilized by aromatic residues (Blue: Y149, F160, and F161) in the Cas7 subunit of Spu Cascade (pale green) through base stacking^45^. Right: The interactions between the protein and the hybridized double strand after replacing Spu Cas7 with Mos350 Cas7 (pink) using Pymol. In both structural diagrams, the left orange strand depicts the DNA strand, and the right orange strand depicts the crRNA strand. **d,** Fold activations of *ASCL1* and *IL1B* genes using Mos350 Cascade-VPR carrying mutations in the Cas7 subunit, compared to wild-type Mos350 Cascade-VPR. **e,** Fold activations of *ASCL1* and *IL1B* genes using three different tools. First: dCas9-VPR; Second: Mos350 Cascade-VPR with L175F and 32 bp spacer (L-F S32); Third: Mos350 Cascade-VPR with L175F and 50 bp spacer (L-F S50). **f,** Fold activations of the *ASCL1* gene and potential off-target genes when targeting the *ASCL1* gene with the three tools in (**e**). The transcription of the *PDCL1P4* gene was slightly decreased when L175F S32 targeted the promoter of *ASCL1* and the transcription of *PRPH* gene was activated when dCas9-VPR targeted the promoter of *ASCL1*. Ctrl or NT: Only the plasmid expressing the Cascade-VPR or dCas9-VPR was transformed. All values are represented as mean ± s.d. of three biological replicates. Statistical significance was assessed by unpaired two-tailed Student’s t-test (ns, not significant; *P < 0.05; **P < 0.01; ***P < 0.001).

Subsequently, we sought to increase the activity by introducing specific amino acid mutations based on protein structure information. Compared to the Spu Cascade subunits, the Mos350 Cascade subunits have similar three-dimensional structures predicted by AlphaFold (Supplementary Fig. 7). In the DNA-bound structure of the Spu Cascade complex, the aromatic residues emanating from the Cas7 thumb (Y149, F160, and F161) form a stacking force that stabilizes the interaction between the crRNA and the target protospacer^45^. The results of sequence alignment and structure prediction showed that the Cas7 of Mos350 I-F2 possesses only one aromatic residue at the corresponding positions (L154, L175, and F176) (Fig. 5c and Supplementary Fig. 8). To increase the DNA-binding activity of the Mos350 Cascade-VPR, we introduced L175F into the Mos350 Cas7 protein, named as L175F-Cascade-VPR. As expected, when targeting the *ASCL1* and *IL1B* loci, L175F-Cascade-VPR exhibited improved activities of up to 2.4-fold and 4.0-fold, respectively, compared to the wild-type Mos350 Cascade-VPR. However, adding the L154Y mutation to L175F-Cascade-VPR did not further enhance transcriptional activation (Fig. 5d).

We combined the two strategies for further improvement, including extending the spacer length to 50 nt and introducing the L175F mutation in Cas7. The activity of the L175F-Cascade-VPR improved to 16.7-fold and 5.3-fold at the *ASCL1* and *IL1B* loci, respectively, after extending the spacer length to 50 nt (Fig. 5e). We also compared the activation activity with that of the dCas9-VPR (SpCas9). The engineered Mos350 Cascade-VPR and the dCas9-VPR efficiently activated gene expression, with the engineered Mos350 performing even better at the *ASCL1* locus (Fig. 5e).

Given the critical importance of binding specificity for transcriptional activators, we conducted a Cascade-dependent DNA off-target analysis of the L175F-Cascade-VPR. Previous research on several type I systems has demonstrated that mismatches in the PAM-proximal region have greater impacts on the activity of CRISPR systems than those in the PAM-distal region, especially those in the region downstream of the position 24^17, 19, 25, 27^, counting the nucleotide adjacent to the PAM as position 1 of the protospacer. Additionally, structural features contribute to the mismatch tolerance at every sixth position of the guide sequence, which flips out of the RNA-DNA heteroduplex structure^45^. Therefore, we searched potential off-target sites with ≤4 mismatches to the crRNA except for mismatches in positions 6, 12, 18, 24−32, located on the promoter regions (≤2 kb upstream of the TSS). The target site was selected at the *ASCL1* locus, which had overlapping target regions of the Mos350 L175F-Cascade-VPR and dCas9-VPR. Its predicted four off-target sites were measured, which also had ≤4 mismatches to SpCas9 crRNAs (Supplementary Fig. 9). For the Mos350 L175F-Cascade-VPR, off-target activation activity was not observed at any of the four potential off-target sites. Importantly, no off-target activation occurred when the spacer length was extended to 50 nt either. However, the dCas9-VPR exhibited activation activity at an off-target site in the *PRPH* promoter region (Fig. 5f). These results indicate that the engineered Mos350 L175F-Cascade-VPR possesses robust target gene activation activity with high specificity.

### I-F2 Cascade-ABE with a wide bimodal editing window

Since the R-loop formed by Cascade is considerably wider than that by Cas9 or Cas12, we investigated the potential of repurposing the Mos350 Cascade as a base editor. As TadA-8e facilitates rapid and efficient deoxyadenosine deamination and is compatible with various Cas proteins^55^, we created four ABEs named 5CABE, 5NABE, 7CABE, and 7NABE by attaching TadA-8e to either the C- or N-terminus of Cas5 or Cas7 in the Mos350 Cascade (L175F) (Fig. 6a). The base editing window was evaluated by targeting a region within the *NIBAN* gene, featuring an alternating 5′-A-N-A-N-3′ sequence^56^. The region could be targeted by either of two crRNAs, resulting in adenines positioned nearly at every even (site 1) or odd (site 2) position, collectively covering positions 4–27 and 30–32 within the protospacer (Fig. 6b). Remarkably, all four ABEs exhibited A·T-to-G·C base editing efficiency, with wide bimodal base editing windows spanning nearly the entire protospacer region (∼30 nt). When TadA-8e was fused to different termini of Cas proteins, the peak in the PAM-proximal end shifted from position 6 to 13. In contrast, the peak in the PAM-distal end (at position 26 or 27) and the valley (at position 16 or 17) remained consistent. The highest base editing efficiency reaching 30.4%, was achieved at position 27 using 5NABE, in which TadA-8e was fused to the N-terminus of Cas5 (Fig. 6b and Supplementary Fig. 10). The 5NABE was investigated in subsequent experiments.

**Fig. 6:**
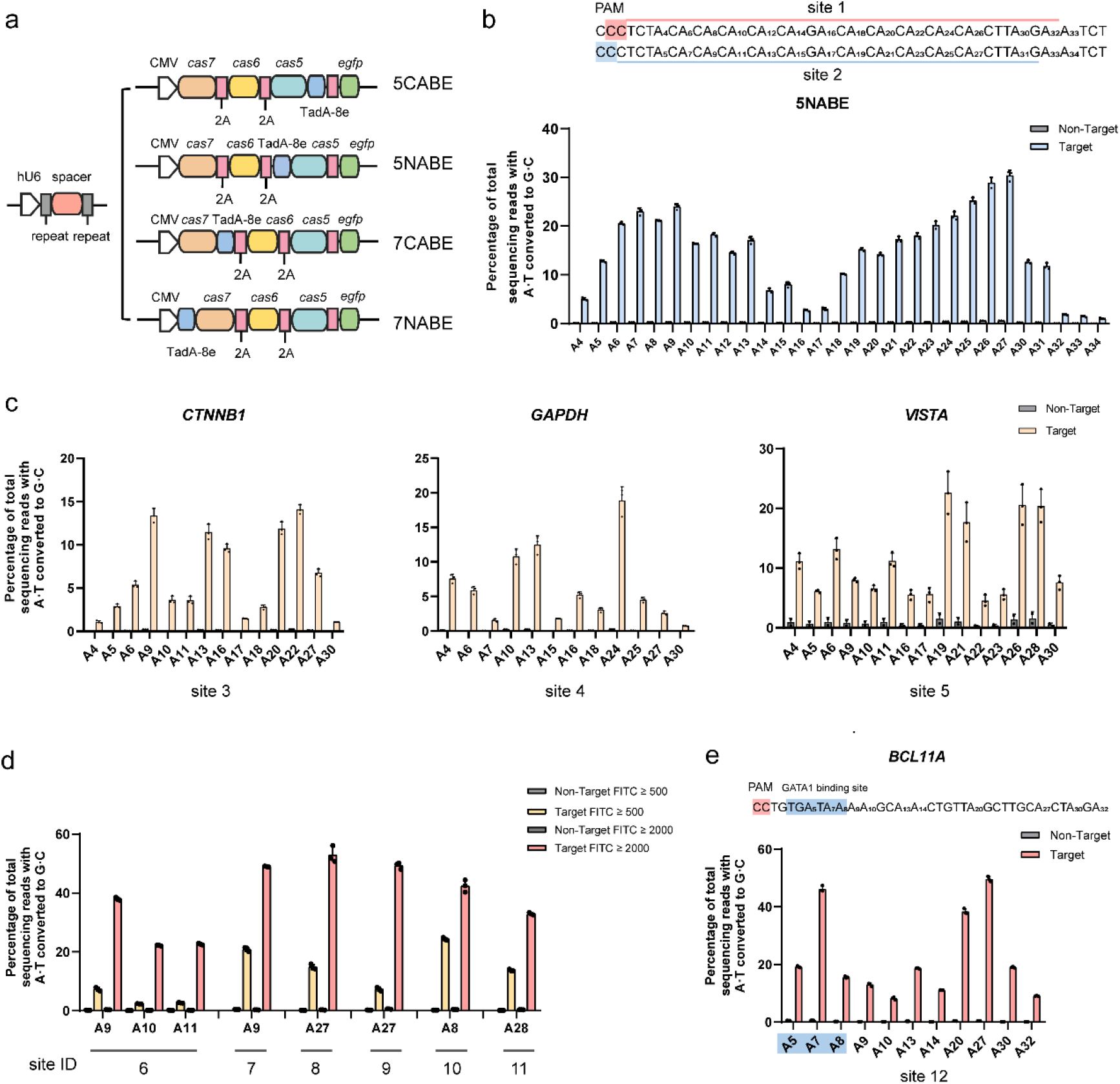
Cascade-ABE8e has a unique wide editing window with a bimodal distribution. **a,** Schematic illustration of gene structures of 5NABE, 5CABE, 7NABE, and 7CABE. TadA-8e was fused to the N-terminus or C-terminus of Cas5 or Cas7. Orange: Cas7; Dark Yellow: Cas6; Cyan: Cas5; Pale blue: TadA-8e; Pink: T2A; Green: EGFP. **b,** The editing window of the 5NABE base editor. The pink area marks the PAM of site 1, and the pink line labels the protospacer. The blue area marks the PAM of site 2, and the blue line labels the protospacer. **c,** A·T-to-G·C editing efficiency of 5NABE at three endogenous (sites 3–5) genomic loci with multiple adenines. **d,** A·T-to-G·C base editing efficiency of 5NABE at six different genomic loci (sites 6–11) in HEK293T cells using different fluorescence thresholds (FITC). **e,** A·T-to-G·C base editing efficiency of 5NABE at the *BCL11A* enhancer GATA1 binding site in HEK293T cells. The pink area marks the PAM of site 12, and the blue area labels the GATA1 binding site. Non-Target: Only the plasmid expressing the Cascade-TadA-8e was transfected. Target: The plasmid expressing the Cascade-ABE and the plasmid expressing the mini-CRISPR were co-transfected. All values in (**b-e**) are represented as mean ± s.d. of three biological replicates, except for non-target results at site 5 of two biological replicates.

We tested the base editing efficiency of 5NABE by targeting different endogenous loci in the genome. We chose four targets (sites 3–6) with multiple adenines and five targets (sites 7–11) with only a single adenine. For targets with multiple adenines, there were varying degrees of A·T-to-G·C conversion in target positions 4–30, with the highest editing efficiency ranging from 7.4% to 22.6%. For targets with a single adenine, significant A·T-to-G·C conversion was observed with the highest editing efficiencies ranging from 7.4% to 24.2% (Fig. 6c and 6d).

We further investigated the applications at disease-related targets. Upregulating fetal hemoglobin expression is a promising therapeutic option for β-thalassemia and sickle-cell disease^57, 58^. *BCL11A* is a transcriptional repressor of fetal hemoglobin. It has been demonstrated that mutations in the core sequence of +58 *BCL11A* erythroid enhancer, especially at the GATA1 binding site, resulted in reduced *BCL11A* expression and concomitant induction of fetal hemoglobin^59–61^. We designed two crRNAs (targeting sites 12 and 13) to install A·T-to-G·C conversion in the core sequence of +58 *BCL11A* erythroid enhancer, with the adenines of the GATA1 binding site located around the two peaks (position 9 or position 27) of the editing window, respectively. The three adenines in GATA1 were edited significantly by the two crRNAs, with a maximum editing efficiency of 46.2% ± 1.1% at position 7 of site 12 (Fig. 6e and Supplementary Fig. 11 and 12). Besides, multi-adenines outside the GATA1 binding site, which also located in the core sequence, were efficiently edited, with a maximum efficiency of 49.6% ± 0.9% at position 27 of site 12 (Fig. 6e and Supplementary Fig. 11 and 12).

Next, we assessed the off-target activity of 5NABE. Potential Cascade-dependent off-target sites were selected with ≤3 mismatches at all positions except mismatches in positions 6, 12, 18, 24–32 of the protospacer. Two targets (sites 5 and 10) and seven corresponding potential off-target sites were selected. Most predicted off-target positions exhibited low off-target activity (Fig. 7). The highest off-target base editing efficiency reaching 4.2%, was achieved at OT2 for site 5. The only two mismatches of OT2 were in positions 22 and 23 in the PAM-distal end, which might be the reason for its high editing efficiency (Fig. 7).

**Fig. 7:**
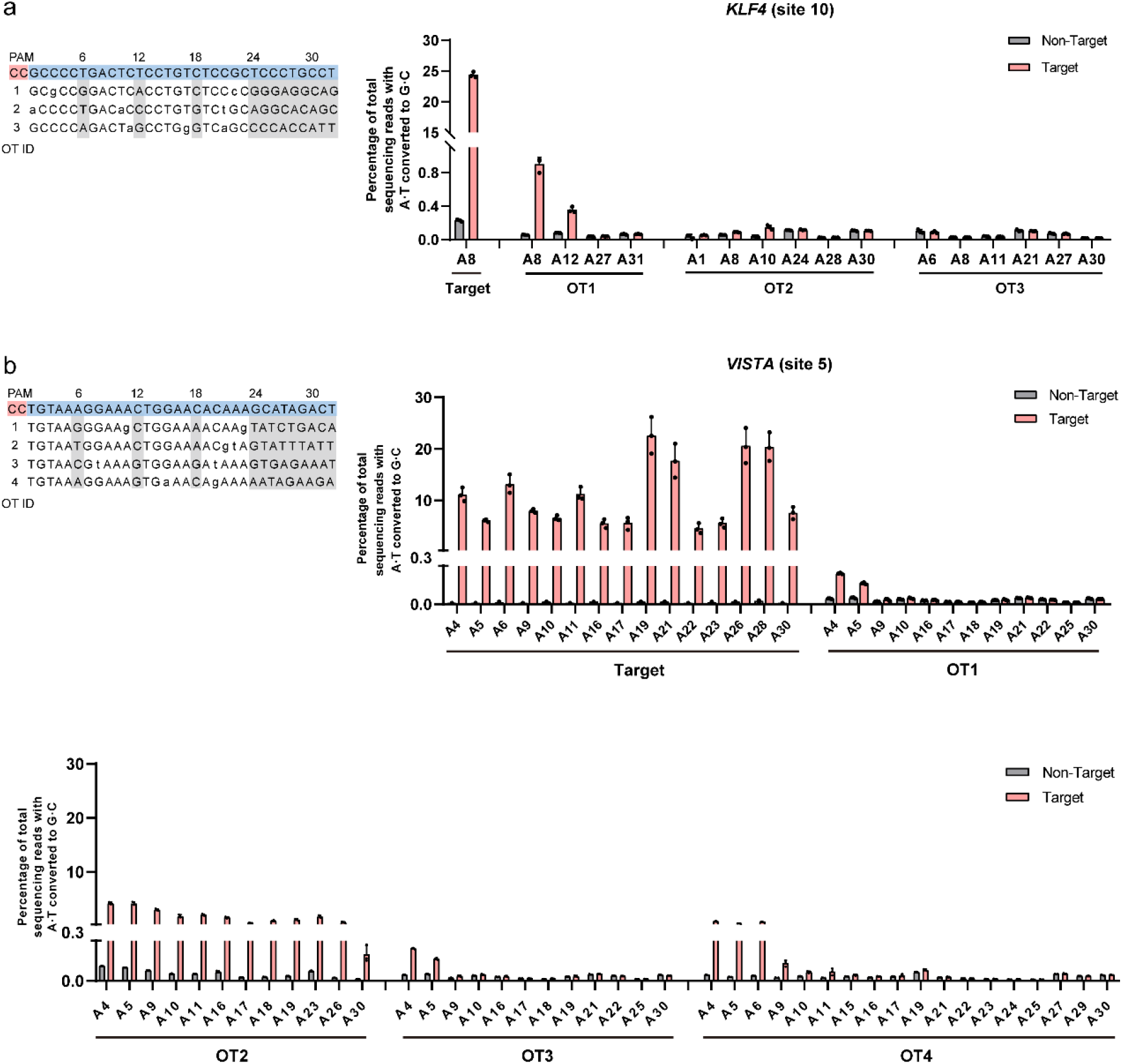
Off-target effects of 5NABE. **a,** Left: Potential Cascade-dependent off-target sites of 5NABE when targeting the *KLF4* gene in HEK293T cells. Lowercase letters represent mismatched bases compared to the target sequence. Right: Base editing efficiency of 5NABE at target (Fig. 6d) and off-target sites. **b,** Left: Potential Cascade-dependent off-target sites of 5NABE when targeting the *VISTA* gene in HEK293T cells. Lowercase letters represent mismatched bases compared to the target sequence. Right: Base editing efficiency of 5NABE at target (Fig. 6c) and off-target sites. The efficiency results of target sites were from Fig. 6. Non-Target: Only the plasmid expressing the Cascade-TadA-8e was transfected. Target: The plasmid expressing the Cascade-ABE and the plasmid expressing the mini-CRISPR were co-transfected. All values were mean ± s.d. of three biological replicates, except for non-target results at site 5 of two biological replicates.

Interestingly, when we raised the fluorescence threshold for EGFP-positive cells in fluorescence-activated cell sorting (FACS) (Supplementary Fig. 13), we found that the editing efficiency was significantly improved. Since EGFP was linked by the self-cleaving 2A peptide to the I-F2 ABE system, the results suggesting that the editing efficiency could be improved by increasing the gene expression level of I-F2 ABE system. Thus, we selected sites 6–11 to test the editing efficiency after raising the fluorescence threshold from 500 to 2000. When cells with stronger fluorescence signal were collected, the A·T-to-G·C conversion efficiencies were significantly improved, with the highest editing efficiencies ranging from 32.6% to 50.8%. Notably, the editing efficiencies for three sites reached approximately 50%, the theoretical maximum editing efficiency (without nickase activity)^36^ (Fig. 6d). These results revealed that the I-F2 ABE system could be very efficient. We also tested the editing efficiency of the predicted off-target sites for site 10 with the fluorescence threshold of 2000. The editing efficiency of position A8 of OT1 was improved from 1.0% to 5.0% (Fig. 7a and Supplementary Fig. 14). These results suggested that the higher editing efficiency might be accompanied with higher off-target activity. If low off-target activity is required, the protein expression level can be reduced or the delivery of protein–RNA complex can be employed^55, 62^.

## Discussion

In this study, we tested 11 type I-F2 Cascades and obtained one system derived from the Mos350 strain, which can be repurposed as a transcriptional activator or base editor in human cells by fusing different domains. The Mos350 Cascade prefers a simple 5′-CC PAM and has a total gene size of ∼2.7 kb, which is significantly smaller than SpCas9 (∼4.1 kb). The pre-crRNA with one spacer flanked by two repeats is 88 bp in length. Multiplexed targeting can be achieved by constructing a CRISPR array containing tandem spacers linked by repeats, which can be processed by Cas6 to target different sites simultaneously. Since the size-minimized AAV backbone for ABE delivery has been developed with small-Cas orthologues like Nme2Cas9 (3.24 kb)^31^, delivering the complete I-F2 Cascade-ABE using a single AAV vector is feasible. Moreover, recent research has found that type I Cascades fused with HNH nuclease domains enable precise dsDNA cleavage, with a total gene size of over 3.3 kb^63^. Compact genome editing tools may be constructed by fusing the type I-F2 Cascade with an HNH domain. Multi-subunit effector complexes of type I systems provide a broad and flexible platform for integrating different functional domains. First, functional components can be fused to various subunits individually or simultaneously across multiple subunits to improve their effects^17, 27–29^. Second, the Cascade complex is very flexible. Modifying the spacer length can alter the Cas protein stoichiometry within the Cascade complex and influence the extent of the R-loop region, potentially impacting its functionality^16, 27, 53, 54, 64^. For transcriptional activation, significant activation was detected with VPR fused to Cas7 (Fig. 4a), which could be attributed to the multi-copy form of Cas7-VPR in Cascade. Moreover, we engineered spacer lengths ranging from −6 nt to +24 nt, resulting in varying degrees of target gene expression (Fig. 5b). This inherent flexibility of the type I system enables delicate regulation of target genes, which is useful for investigating gene functions and biological mechanisms. However, for base editing, the highest editing efficiency was obtained with TadA-8e fused to Cas5 (Fig. 6b and Supplementary Fig. 10). Besides, in immunofluorescence analysis, we could only detect the signals of all three Cas proteins in partial cells. Thus, higher activation efficiency may be achieved by optimizing the NLS.

Current base editors mainly contain catalytically impaired Cas9 or Cas12 orthologs fused to enzymes that alter the DNA bases, which can help study the effects of mutations within genes and treat genetic diseases^36, 38, 39, 55^. The editing window is limited by the R-loop size of Cas9 or Cas12, which is hard to expand over 20 nt^40, 65^. However, the potential of base editing with Cascade, which induces a large R-loop, has not yet been explored. We constructed ABEs by fusing the I-F2 Cascade with TadA-8e. The editing window with a bimodal distribution covered almost the entire R-loop region, from positions 4 to 31 (Fig. 6b and Supplementary Fig. 10), whereas the Cas9- or Cas12-derived ABE8e typically had a unimodal window of less than 12 nt^55^. The wide editing window can expand the range of targetable sites and is useful for disruption of functional sequences, like disruption of infection-related genes or inhibitory genes for genetic therapy. The wide editing window is also useful for in situ saturating mutagenesis of important genes, and multiplex base editing. For example, expanding the editing window is an important aspect for crop breeding^65^.

On the other hand, among the four ABEs we created, 5NABE is the most efficient design. With 5NABE, the editing efficiency for several targets could be close to 50%, the theoretical maximum editing efficiency (without nickase activity)^36^ (Fig. 6d). It is also more efficient than most ABEs based on dCas12, such as enAsABE8e derived from dead engineered AsCas12a^55^, dCasMINI-ABE derived from dead engineered Cas12f^39^, and dCas12Pro-ABE derived from dead Cas12n^66^. Besides, without the nickase activity, the editor may be safer for application^67^.

Off-target activation efficiency was not observed for Mos350 L175F-Cascade-VPR, even with an extended spacer of 50 nt. In contrast, a significant off-target site was detected for SpCas9 (Fig. 5f). For base editing, with the increase of protein expression, the editing efficiency of target sites improved significantly (Fig. 6d). It should be noted that higher off-target editing efficiency may be induced at the same time (Supplementary Fig. 14). There are multiple factors that determine the editing efficiency of off-target sites, like the number, location, and type of mismatches^68^. Large numbers of gRNAs need to be tested to understand the specificity of the system. In addition, off-target efficiency can be reduced by engineering proteins^55^ and gRNAs^69^, as well as delivery of protein–RNA complex^55, 62^.

In summary, we demonstrate that the miniaturized I-F2 Cascade, which is significantly smaller than SpCas9, can be developed into programmable tools for use in human cells. By engineering the crRNA and Cascade, the activation efficiency of the Cascade-VPR can match or surpass that of the dCas9-VPR. Moreover, the inherent flexibility of the type I system enables delicate regulation of target genes. We also explored the potential of base editing with type I Cascade and successfully created an ABE with type I-F2 Cascade (5NABE). The 5NABE induces a significantly wide editing window of ∼30 nt, which can be useful for genetic screening and disrupting the functional sequences. In addition, systematic engineering of the crRNA and Cascade components and optimizing the architecture and NLS may further improve the on-target efficiency and reduce the off-target activity.

## Methods

### Identification of type I-F2 systems

We downloaded genomic data of bacteria and archaea from NCBI in 2020, constructing a representative dataset by selecting one genome per species. Additionally, we obtained human gut metagenomic data from several articles^70–72^. CRISPR array was predicted by MinCED (version 0.2.0) software. Cas7, Cas5, and Cas6 proteins were identified by the following three methods with HMMER (version 3.3.2) software: 1. Using Pfam domain (Cas7: PF05107, Cas6: PF09559, PF10040, PF01881, PF17262, PF17952, PF17955, PF19308, Cas5: PF09704, e-value cutoff of 1E-5); 2. Using the previously described CRISPR-Cas protein^1, 73^; 3. From the CRISPRCasdb database^74^. To find out more I-F2 systems, profiles of I-F2 Cas5/6/7 sequences were generated using MAFFT (version 7.487)^75^ and hmmbuild software. Proteins from constructed representative dataset and metagenome were searched against the HMM profiles using hmmsearch. The cluster simultaneously containing proteins Cas5, Cas6, and Cas7 of I-F2 was selected as a candidate I-F2 system.

### Phylogenetic tree analysis

After being manually confirmed, 507 Cas7 proteins from different subtypes were aligned using MAFFT (version 7.487) software. The maximum likelihood (ML) phylogenetic tree was constructed using IQ-TREE (version 2.0.3) software^76^, with automatic model selection and 1000 bootstraps and then visualized by the iTOL website. The protein sequences of Cas7 used to construct the phylogenetic tree are listed in Supplementary Table 1. For the phylogenetic tree analysis of the I-F2 systems shown in Supplementary Fig. 1, we concatenated Cas7, Cas5, and Cas6 from I-F2 CRISPR-Cas systems and constructed the phylogenetic tree with the same method. Multiple sequence alignment of the repeats was conducted using Geneious Prime (version 2023.2.1).

### Plasmid construction

For the expression of the I-F2 CRISPR-Cas system in *E. coli*, *cas5-7* from eight candidate systems were *E. coli* codon-optimized, synthesized (GeneScript), and ligated to the pETDuet vector (predigested with NdeI and KpnI) using T4 DNA ligase. The mini-CRISPR fragments derived from the primer annealing extensions were ligated to the pETDuet vector (predigested with XbaI and BamHI) using T4 DNA ligase. The Mos350 *cas3* gene was *E. coli* codon-optimized and synthesized into the pCOLADuet vector within NdeI and KpnI (GeneScript).

For transcriptional activation in human cells, *dcas9*, *cas5-7* from 11 candidate systems and VPR transcriptional activator^27^ were human codon-optimized, synthesized (GeneScript), and integrated into the pcDNA3.1 vector (predigested with XhoI and KpnI) by Gibson assembly. For base editing in human cells, the protein-coding sequences and the linearized pcDNA3.1-P2A-EGFP (predigested with XbaI and KpnI) were ligated by Gibson assembly. Mini-CRISPRs (repeat-BbsI-BbsI-repeat) and the U6 promoter were inserted into the linearized pcDNA3.1 vector (predigested with MluI and BbsI) by Gibson assembly. Oligonucleotides encoding spacers were annealed and ligated into linearized vectors (predigested with BbsI) using T_4_ DNA ligase. The Gibson assembly was conducted using the Hieff Clone® Plus Multi One Step Cloning Kit (YESEN). The restriction enzymes and T4 DNA ligase were ordered from New England BioLabs. The full sequences of the plasmids used in the paper are listed in Supplementary Table 1.

### Measurement of bacterial growth and fluorescence

For fluorescence measurements, the plasmid expressing the Cascade and mini-CRISPR and the plasmid expressing the YFP reporter were co-transformed into *E. coli* BL21 (DE3). The transformant was inoculated into 3 mL of Luria-Bertani (LB) medium with antibiotics (100 µg/mL carbenicillin, 34 µg/mL chloramphenicol) and grown overnight at 200 r.p.m. and 37°C. Then, the seed broth was inoculated at 1:100 into M9 medium (1 × M9 salt, 0.1 mM CaCl_2_, 2 mM MgSO_4_, 10 g/mL thiamine hydrochloride, and 1% (w/v) tyrosine acid) containing antibiotics of the same concentrations and 0.05 mM isopropyl-β-D-thiogalactoside (IPTG). After cultivation at 30°C for 12 h, the OD_600_ and fluorescence values were measured using the Synergy H4 Hybrid Reader (BioTek). The fluorescence measurement wavelength is set as follows: 500 nm for YFP excitation, 550 nm for YFP emission. Fluorescence values were normalized to OD and the relative YFP fluorescence values of cultures were calculated as the ratio between cultures induced by 50 μM IPTG and cultures induced by 0 μM IPTG.

### PAM library construction

The protospacer with four randomized nucleotides upstream (5′ end) was achieved by PCR with primers PRL1 and PRL2 using the pTemplate vector as the template and ligated to the linearized plasmid pACYC184 fragment (PCR by primers PRL3 and PRL4) by Gibson assembly. The DH5α-competent cells were transformed with the ligation products. More than 10^4^ cells were collected, and library plasmids were extracted using the TIANpure Midi Plasmid Kit (TIANGEN).

### PAM depletion assay

The plasmid expressing the Cascade and mini-CRISPR and the plasmid expressing Cas3 were co-transformed into *E. coli* BL21 (DE3) cells. Subsequently, electrocompetent cells were prepared through ice-cold H_2_0 and 10% glycerol washing. The competent cells were electroporated with 200 ng of library plasmids containing four randomized nucleotides at the upstream (5′ end) of the target sequence. After 2 h of recovery, 10 μL of the culture was inoculated into 1 mL of LB and spread on the selective medium, and the colony-forming units were measured to ensure proper coverage of all possible combinations of four randomized nucleotides. Recovery cultures were transferred to 10 mL of liquid medium containing appropriate antibiotics (100 µg/mL carbenicillin, 34 µg/mL chloramphenicol, and 50 µg/mL kanamycin) and 0.05 mM IPTG (or 0 mM IPTG as a negative control), which ensured plasmid propagation and Cascade-Cas3 effector production, and grew at 25°C for 48 h. The propagated plasmids were extracted using the TIANpure Midi Plasmid Kit (TIANGEN) and analyzed relative to the non-induced control. The target region was amplified by PCR with primers containing different barcodes and deep-sequenced with the Illumina NovoSeq 6000 platform with the PE150 strategy. Sequencing reads were aligned to the corresponding plasmids and PAM randomized regions were extracted. The abundance of each possible four-nucleotide combination was counted from the mapped reads and normalized to the total reads for each sample. The enrichment score of each PAM was the ratio compared to the abundance in the control group without IPTG induction. PAMs with enrichment scores greater than 2.5 were selected and visualized by WebLogo^77, 78^.

### Plasmid interference assay

The plasmid expressing the Cascade and mini-CRISPR (the plasmid expressing the Cascade without the mini-CRISPR was used in the negative control) and plasmids expressing Cas3 were co-transformed into *E. coli* BL21 (DE3). Each transformant was inoculated into 3 mL of LB medium containing antibiotics (100 µg/mL carbenicillin, 34 µg/mL chloramphenicol, and 50 µg/mL kanamycin), cultured overnight and then inoculated into 50 mL of LB medium at a ratio of 1:100 with the same antibiotics concentrations. Cultures were grown to an OD_600_ of 0.3 and then added with different concentrations of IPTG (0.05 mM, 0.1 mM and 0.2 mM) until the OD_600_ was between 0.6 and 0.7. Electrocompetent cells were prepared by washing with ice-cold H_2_0 and 10% glycerol. The target plasmid (100 ng) was electroporated. The LB medium (1mL) was added to recover the culture for 1 h. A series of gradient dilutions were performed, and 5 µL was added dropwise to the LB plate with antibiotics (100 µg/mL carbenicillin, 34 µg/mL chloramphenicol, and 50 µg/mL kanamycin). The plates were observed after overnight incubation at 37℃.

### Cell culture and transfection

HEK293T cells were grown in Dulbecco’s Modified Eagle’s Medium (Invitrogen) with 10% Fetal Bovine Serum (FBS) and 1% penicillin-streptomycin, and the cells were cultured at 37°C in an incubator with 5% CO_2_. A medium (1 ml) containing 3 × 10^5^ cells was added to each well of a 12-well plate. When the cell fusion was 70–85% (∼24 h), 1.2 µg of the Cascade expression plasmid, 0.8 µg of the mini-CRISPR expression plasmid and 2 μL of PEI were added to 200 μL of DMEM medium, and the mixture was placed at room temperature for 15 min. In a comparison control, only 1.2 µg of the Cascade expression plasmid or no plasmid was transfected, respectively. Then, the mixture was added with 550 μL of 2% FBS DMEM medium and mixed well. The medium in the well plate was replaced with the mixture. After 6 h, the mixture was replaced with 10% FBS DMEM medium.

### Immunofluorescence staining

HEK293T cells were transfected with 1.2 µg of the Cascade expression plasmid and 0.8 µg of the mini-CRISPR expression plasmid in 12-well plates. After 1 d, the cells were digested, and 5 × 10^4^ cells were added to a 35 mm glass bottom. After incubation for 1 d, the cells were washed with PBS and fixed in 4% paraformaldehyde. Cells were incubated with blocking solution (1% BSA PBS, 0.1% Triton X-100 PBS) and then incubated with mouse anti-Flag RBITC (1: 300 dilution, Bioss, bs-33346M-RBITC), rabbit anti-Myc FITC (1: 300 dilution, Bioss, bs-0842R-FITC), and rabbit anti-HA AF647 (1: 300 dilution, Bioss, bs-0966R-A647). The cells were then incubated with DAPI for nucleic acid staining and imaged using a SP8 fluorescence microscope (Leica).

### Co-immunoprecipitation and Western blot

HEK293T cells were co-transfected with 2.4 μg of the Cascade expression plasmid and 1.6 μg of the crRNA expression plasmid in 6-well plates. After 1 d, the cells were lysed with 500 uL of the Cell lysis buffer for Western and IP (Beyotime) on ice for 30 min, with protease inhibitor cocktail (Roche) and 1 mM phenylmethanesulfonylfluoride added. Samples were centrifuged at 13523 rcf for 5 min at 4°C. For co-immunoprecipitation analysis, 400 uL of the supernatant were immunoprecipitated using mouse anti-flag-agarose conjugate (Sigma-Aldrich) at 4°C for 3 h. The immunoprecipitation products were washed four times with IP Lysis Buffer, then mixed with 5 × loading buffer (10% SDS, 0.5% bromophenol blue, 50% glycerol, 5% β-mercaptoethanol, 0.25 M pH 6.8 Tris-HCl), and heated at 100°C for 10 min as output. The rest 100 uL of the supernatant were mixed with 5 × loading buffer and heated at 100°C for 10 min as input. Samples were loaded onto 12% SDS-PAGE and electrophoresed for 1.5 h at 100 V in the running buffer (0.1% SDS, 25 mM Tris, 192 mM glycine). Proteins were transferred to PVDF membranes in the transfer buffer (20% methanol, 25 mM Tris, 190 mM Glycine) at 300 mA for 1.25 h at 4°C. The blot was blocked with 5% milk-TBST (10 mM Tris, 150 mM NaCl, and 0.1% Tween-20) for 2 h at room temperature, followed by incubation with mouse anti-Flag (1:5000 dilution, Sigma-Aldrich, M2 clone F3165) or rabbit anti-Myc (1:5000 dilution, CapitalBio, P1007t) in 5% milk-TBST at 4°C overnight. Then the blots were washed with TBST, followed by incubation with goat anti-mouse conjugated horseradish peroxidase (HRP) (1:5000 dilution, Zhongshan Golden Bridge, ZB-2305) or goat anti-rabbit conjugated HRP (1:5000 dilution, Zhongshan Golden Bridge, ZB-2301) in 5% milk-TBST for 2 h at room temperature. The blots were washed in TBST and then visualized using Western-ECL (Thermo Fisher) substrates on a Tanon 5200 Multi Chemiluminescent Imaging System (Tanon).

### RNA analysis

The Cascade-VPR expression plasmid (1.2 μg) and mini-CRISPR expression plasmid (0.8 μg) were co-transfected into HEK293T cells in 12-well plates. Total RNA was extracted using a SteadyPure Universal RNA Extraction Kit II (Accurate Biology) after 2 d. For qPCR analysis, the total RNA of each sample (1.25 μg) was used for reverse transcription in a 20 μL reaction using the ABScript III RT Master Mix for qPCR with a gDNA Remover kit (Abclonal). In each reaction, 2 μL of the cDNA was used with the KAPA SYBR FAST Universal qPCR Kit (KAPA) and run on a CFX96 real-time PCR detection system (Bio-Rad). The qPCR primers are listed in Supplementary Table 1. All qPCR data were expressed as fold change in RNA, normalized to GAPDH expression, and analyzed relative to the control group. The relative expression level was determined by the 2^−ΔΔCt^ method^79, 80^. To normalize data from different TSS between genes (*ACTC1*, *ASCL1*, and *NEUROD1*), data in the same gene were normalized by percentage, and 100% was defined by the sum of all values. The target sites were divided into three regions based on the distance to the TSS (0−250 bp, 250−500 bp, >500 bp), and the three sites with the highest percentage in each of the three regions were picked out for each target gene. Normalized values for all 3 genes are plotted as box-and-whisker plots with min to max as an option.

For Northern blot analysis, the total RNA of each sample (10 μg) and the Century-Plus RNA ladder (Thermo Fisher Scientific) were mixed with an equal volume of RNA loading dye (New England Biolabs), respectively, and denatured in a water bath at 65°C for 10 min, and placed on ice for 2 min. Samples were centrifuged, loaded onto 8% polyacrylamide gel, and electrophoresed at 200 V for 40 min in TBE buffer. The lane including the RNA marker was excised, stained with ethidium bromide, and imaged using a GenoSens2100 System (CLINX). The RNA samples were transferred onto a Biodyne B nylon membrane (Pall) in TBE buffer using the Mini-Protean Tetra system (Bio-Rad) and then the membrane was UV cross-linked. The hybridization signal from the biotin-labeled probe (listed in Supplementary Table 1) was detected using the Chemiluminescent Nucleic Acid Detection Module Kit (Thermo Fisher Scientific), following the manufacturer’s protocol. The membrane was imaged using a Tanon 5200 Multi chemiluminescent imaging system (Tanon).

For small RNA sequencing, the total RNA of each sample (50 μg) was treated with 1 μL of polynucleotide kinase (New England Biolabs) in 50 μL of total solution at 37℃ for 6 h, followed by incubation at 37℃ for 2 h with 5 μL of 10 mM ATP. Then, the proteins were removed using the phenol-chloroform method. Purified RNA was treated with RNA 5′ pyrophosphohydrolase (New England Biolabs) for 2 h at 37℃. RNA was purified by phenol-chloroform extraction for sequencing. RNA molecules ranging from 50 to 100 nt were selected to construct a small RNA library using the NEXTFLEX Small RNA-Seq Kit (Bioo Scientific) and then subjected to Illumina HiSeq sequencing (paired-end, 150-bp reads). The raw data was processed to remove the adapters using Cutadapt (version 3.5) software^81^. The resulting reads were mapped to the mini-CRISPR sequence using the BLASTN software, and the coverage depth of each site was calculated using custom Perl scripts.

### Structure analysis

The superposition of three-dimensional structures of Cas proteins were analyzed using PyMol (version 2.5.5) software. The structure of SpuCascade^45^ (5O6U, https://doi.org/10.2210/pdb5O6U/pdb) was downloaded from the PDB database. The structure of Mos350 Cas5 (A0A378Q7F6), Mos350 Cas6 (A0A378Q7P1), and Mos350 Cas7 (A0A378Q7G2) were downloaded from AlphaFoldDB^82, 83^. Sequence alignment of Cas7 proteins from SpuCN32 and Mos350 was performed by MUSCLE alignment using Geneious Prime (version 2023.2.1).

### Off-target analysis

To predict potential off-target sites of the dCas9-VPR, we searched for sites of ≤4 mismatches with SpCas9 sgRNA. To predict Mos350 Cascade-VPR off-target sites, disregarding the mismatches at positions 6, 12, 18, and 24–32 of the protospacer, we searched for all possible off-targets with ≤4 mismatches to the remaining positions of the protospacer using Cas-OFFinder^84^. All possible off-target sites located in the promoter region of a given gene (≤2 kb upstream of the TSS) are listed in Supplementary Table 1. To predict Mos350 Cascade-TadA8e off-target sites, we searched for sites of ≤3 mismatches, disregarding the mismatches at positions 6, 12, 18, and 24–32 of the protospacer, on the whole genome using Cas-OFFinder^84^. The sequences of all putative off-target sequences are listed in Supplementary Table 1.

### Genomic DNA extraction, high-throughput DNA sequencing, and data analysis

At 72 h post-transfection, cells were washed with PBS. Then cells were digested with 0.25% trypsin (Gibco) for FACS. Samples were performed on a flow cytometer (BD influx) after passing through a 70 μm cell strainer. Linear SSC-A versus linear FSC-A was used to select the live HEK293T cells. Within the gate, FSC-W versus FSC-H and SSC-H versus SSC-W were used to gate on single cells. Then within the single live cell gate, we utilized EGFP-negative control to delineate the boundary between positive and negative cells. The final gate was utilized to sort GPF-positive single live cells. Then, cells that were positive for the EGFP signal (20000–40000 cells) were collected and lysed with 30–60 μL lysis solution (per liter, 10 mL of 1 M Tris-HCl, pH 7.5, 5 mL of 10% (wt/vol) SDS solution, adding nuclease-free water to a final volume of 1 L) with 1:1000 (vol/vol) proteinase K (New England Biolabs) at 37°C for 1 h. After heated at 80°C for 20 min, the samples were used as PCR templates. The primers used for PCR amplification of the target and off-target genomic loci are shown in Supplementary Table 1. PCR products were purified by 2% agarose gel electrophoresis. PCR products with different barcodes were mixed and deep-sequenced using the Illumina NovoSeq 6000 platform. Editing results were analysed using CRISPResso2^85^ with the following parameters: minimum of 60% homology for alignment to the amplicon sequence, quantification window of 20 bp, quantification_window_center of –15. Base editing values are representative of two or three independent biological replicates, with mean ± s.d. shown. The base editing values were calculated as the percentage of reads with A·T-to-G·C conversions to the total aligned reads.

### Statistical analysis

All data analyzed was presented as mean ± s.d. of two or three biological replicates. All P values were calculated by one-way analysis of variance (ANOVA) with Dunnett’s multiple comparisons tests (α = 0.05) (for data having more than two groups) or unpaired two-tailed Student’s t-test (for data having only two groups). Statistical significance level: n.s., not significant; *P < 0.05; **P < 0.01; ***P < 0.001. All P values are listed in Supplementary Table 2.

## Data availability

The data of small RNA sequencing and high-throughput DNA sequencing were deposited in the National Microbiology Data Center (NMDC) database under accession numbers NMDC40043697, NMDC40043744, NMDC40043745, and NMDC40043812. All the other data are available in the main text or supplementary materials.

## Code availability

Custom scripts for Fastq barcode split of samples in base editing analysis and calculating the coverage depth of each site in small RNA sequencing are available in Supplementary Note.

## Supporting information

Custom scripts

## Acknowledgements

H.X. discloses support for the research of this work from the Strategic Priority Research Program of the Chinese Academy of Sciences [XDA24020101], the National Natural Science Foundation of China [32230061], and the National Key R&D Program of China [2020YFA0906800]. L.G. discloses support for the research of this work from the National Natural Science Foundation of China [32100499; 32150020]. We thank Wei Jiang, Xinyu Li, and Dahui Zhao for help with HEK293T cell culture, co-immunoprecipitation, and immunofluorescence assays. We thank Tong Zhao for help with flow cell sorting and Zehua Chen for help with fluorescence reporter system construction in *E. coli*. We thank Pengju Yu for his help in the qPCR experiments. We thank all members of the Xiang laboratory for helpful advice and discussion.

## Author contributions

L.G., J.G., and H.X. designed the project, performed analyses, and wrote the manuscript. J.G. conducted experiments. Z.L., S.F., and C.Y. participated in vector construction and cell experiments. H.Y. performed bioinformatic analyses. M.L., J.H., and D.Z. performed analyses. H.X. and L.G. supervised the research. All the authors have proofread the manuscript.

