## Supplementary Figure for "Engineered minimal type I CRISPR-Cas system for transcriptional activation and base editing in human cells"

**Contents:**

**Supplementary Figures 1–14**

Supplementary Fig. 1 | Phylogenetic analysis of 93 I-F2 CRISPR-Cas systems.
Supplementary Fig. 2 | Analyses of CRISPR arrays next to the I-F2 Cas operon.
Supplementary Fig. 3 | Mos350 I-F2 system provides immunity against plasmids containing protospacers with CC PAM at the 5′ end.
Supplementary Fig. 4 | Source information of the I-F2 CRISPR-Cas systems utilized in the experiments.
Supplementary Fig. 5 | Small RNA sequencing reveals Cas6 cleavage site in repeat sequence.
Supplementary Fig. 6 | Optimizing transcriptional activation efficiency of Cascade-VPR with different linkers and gene arrangements.
Supplementary Fig. 7 | Superposition of three-dimensional structures of Cas5, Cas6, Cas7, and Cascade from Spu and Mos350 I-F2 systems.
Supplementary Fig. 8 | Sequence alignment between Spu Cas7 and Mos350 Cas7.
Supplementary Fig. 9 | Predicted off-target sites of the Mos350 Cascade-VPR and dCas9-VPR targeting the *ASCL1* locus.
Supplementary Fig. 10 | Base editing windows of three different I-F2 ABE constructions in human cells.
Supplementary Fig. 11 | Base editing efficiency of 5NABE at the GATA1 site of the *BCL11A* enhancer.
Supplementary Fig. 12 | Allele compositions following treatment with 5NABE at the GATA1 binding site of the *BCL11A* enhancer.
Supplementary Fig. 13 | FACS gating examples for EGFP-positive cells sorting conditions.
Supplementary Fig 14 | Off-target efficiencies edited by of 5NABE at site 10 with a higher fluorescence threshold.

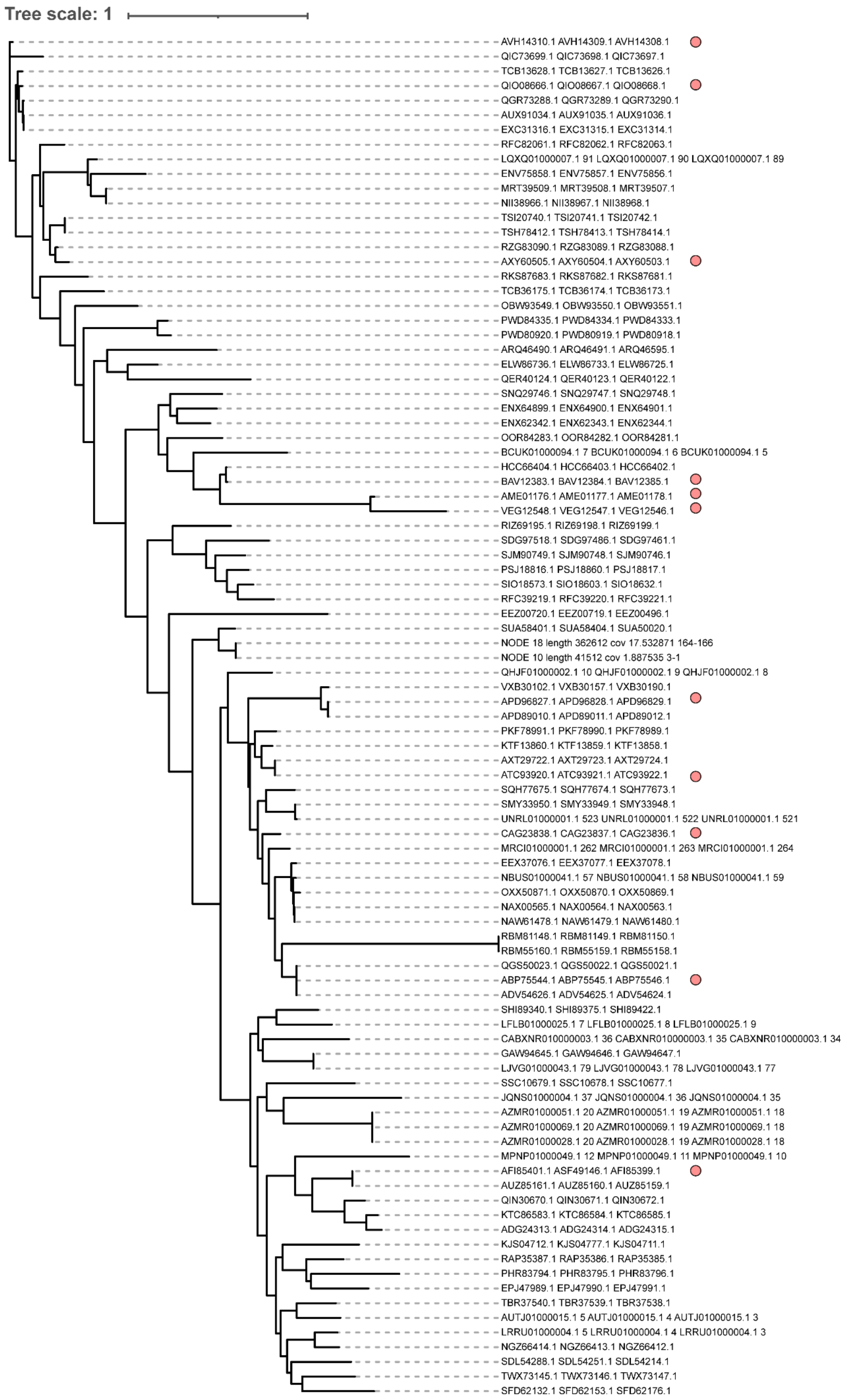

Tree scale: 1
AVH14310.1 AVH14309.1 AVH14308.1
QIC73699.1 QIC73698.1 QIC73697.1
TCB13628.1 TCB13627.1 TCB13626.1
QIO08666.1 QIO08667.1 QIO08668.1
QGR73288.1 QGR73289.1 QGR73290.1
AUX91034.1 AUX91035.1 AUX91036.1
EXC31316.1 EXC31315.1 EXC31314.1
RFC82061.1 RFC82062.1 RFC82063.1
LQXQ01000007.1 91 LQXQ01000007.1 90 LQXQ01000007.1 89
ENV75858.1 ENV75857.1 ENV75856.1
MRT39509.1 MRT39508.1 MRT39507.1
NII38966.1 NII38967.1 NII38968.1
TSI20740.1 TSI20741.1 TSI20742.1
TSH78412.1 TSH78413.1 TSH78414.1
RZG83090.1 RZG83089.1 RZG83088.1
AXY60505.1 AXY60504.1 AXY60503.1
RKS87683.1 RKS87682.1 RKS87681.1
TCB36175.1 TCB36174.1 TCB36173.1
OBW93549.1 OBW93550.1 OBW93551.1
PWD84335.1 PWD84334.1 PWD84333.1
PWD80920.1 PWD80919.1 PWD80918.1
ARQ46490.1 ARQ46491.1 ARQ46595.1
ELW86736.1 ELW86733.1 ELW86725.1
QER40124.1 QER40123.1 QER40122.1
SNQ29746.1 SNQ29747.1 SNQ29748.1
ENX64899.1 ENX64900.1 ENX64901.1
ENX62342.1 ENX62343.1 ENX62344.1
OOR84283.1 OOR84282.1 OOR84281.1
BCUK01000094.1 7 BCUK01000094.1 6 BCUK01000094.1 5
HCC66404.1 HCC66403.1 HCC66402.1
BAV12383.1 BAV12384.1 BAV12385.1
AME01176.1 AME01177.1 AME01178.1
VEG12548.1 VEG12547.1 VEG12546.1
RIZ69195.1 RIZ69198.1 RIZ69199.1
SDG97518.1 SDG97486.1 SDG97461.1
SJM90749.1 SJM90748.1 SJM90746.1
PSJ18816.1 PSJ18860.1 PSJ18817.1
SIO18573.1 SIO18603.1 SIO18632.1
RFC39219.1 RFC39220.1 RFC39221.1
EEZ00720.1 EEZ00719.1 EEZ00496.1
SUA58401.1 SUA58404.1 SUA50020.1
NODE 18 length 362612 cov 17.532871 164-166
NODE 10 length 41512 cov 1.887535 3-1
QHJF01000002.1 10 QHJF01000002.1 9 QHJF01000002.1 8
VXB30102.1 VXB30157.1 VXB30190.1
APD96827.1 APD96828.1 APD96829.1
APD89010.1 APD89011.1 APD89012.1
PKF78991.1 PKF78990.1 PKF78989.1
KTF13860.1 KTF13859.1 KTF13858.1
AXT29722.1 AXT29723.1 AXT29724.1
ATC93920.1 ATC93921.1 ATC93922.1
SQH77675.1 SQH77674.1 SQH77673.1
SMY33950.1 SMY33949.1 SMY33948.1
UNRL01000001.1 523 UNRL01000001.1 522 UNRL01000001.1 521
CAG23838.1 CAG23837.1 CAG23836.1
MRCI01000001.1 262 MRCI01000001.1 263 MRCI01000001.1 264
EEX37076.1 EEX37077.1 EEX37078.1
NBUS01000041.1 57 NBUS01000041.1 58 NBUS01000041.1 59
OXX50871.1 OXX50870.1 OXX50869.1
NAX00565.1 NAX00564.1 NAX00563.1
NAW61478.1 NAW61479.1 NAW61480.1
RBM81148.1 RBM81149.1 RBM81150.1
RBM55160.1 RBM55159.1 RBM55158.1
QGS50023.1 QGS50022.1 QGS50021.1
ABP75544.1 ABP75545.1 ABP75546.1
ADV54626.1 ADV54625.1 ADV54624.1
SHI89340.1 SHI89375.1 SHI89422.1
LFLB01000025.1 7 LFLB01000025.1 8 LFLB01000025.1 9
CABXNR010000003.1 36 CABXNR010000003.1 35 CABXNR010000003.1 34
GAW94645.1 GAW94646.1 GAW94647.1
LJVG01000043.1 79 LJVG01000043.1 78 LJVG01000043.1 77
SSC10679.1 SSC10678.1 SSC10677.1
JQNS01000004.1 37 JQNS01000004.1 36 JQNS01000004.1 35
AZMR01000051.1 20 AZMR01000051.1 19 AZMR01000051.1 18
AZMR01000069.1 20 AZMR01000069.1 19 AZMR01000069.1 18
AZMR01000028.1 20 AZMR01000028.1 19 AZMR01000028.1 18
MPNP01000049.1 12 MPNP01000049.1 11 MPNP01000049.1 10
AFI85401.1 ASF49146.1 AFI85399.1
AUZ85161.1 AUZ85160.1 AUZ85159.1
QIN30670.1 QIN30671.1 QIN30672.1
KTC86583.1 KTC86584.1 KTC86585.1
ADG24313.1 ADG24314.1 ADG24315.1
KJS04712.1 KJS04777.1 KJS04711.1
RAP35387.1 RAP35386.1 RAP35385.1
PHR83794.1 PHR83795.1 PHR83796.1
EPJ47989.1 EPJ47990.1 EPJ47991.1
TBR37540.1 TBR37539.1 TBR37538.1
AUTJ01000015.1 5 AUTJ01000015.1 4 AUTJ01000015.1 3
LRRU01000004.1 5 LRRU01000004.1 4 LRRU01000004.1 3
NGZ66414.1 NGZ66413.1 NGZ66412.1
SDL54288.1 SDL54251.1 SDL54214.1
TWX73145.1 TWX73146.1 TWX73147.1
SFD62132.1 SFD62153.1 SFD62176.1

**Supplementary Fig. 1 | Phylogenetic analysis of 93 I-F2 CRISPR-Cas systems.** The protein sequences of Cas7-Cas5-Cas6 were concatenated and used to construct an evolutionary tree using the maximum likelihood method. 11 systems marked by pink circles were randomly selected for the research.

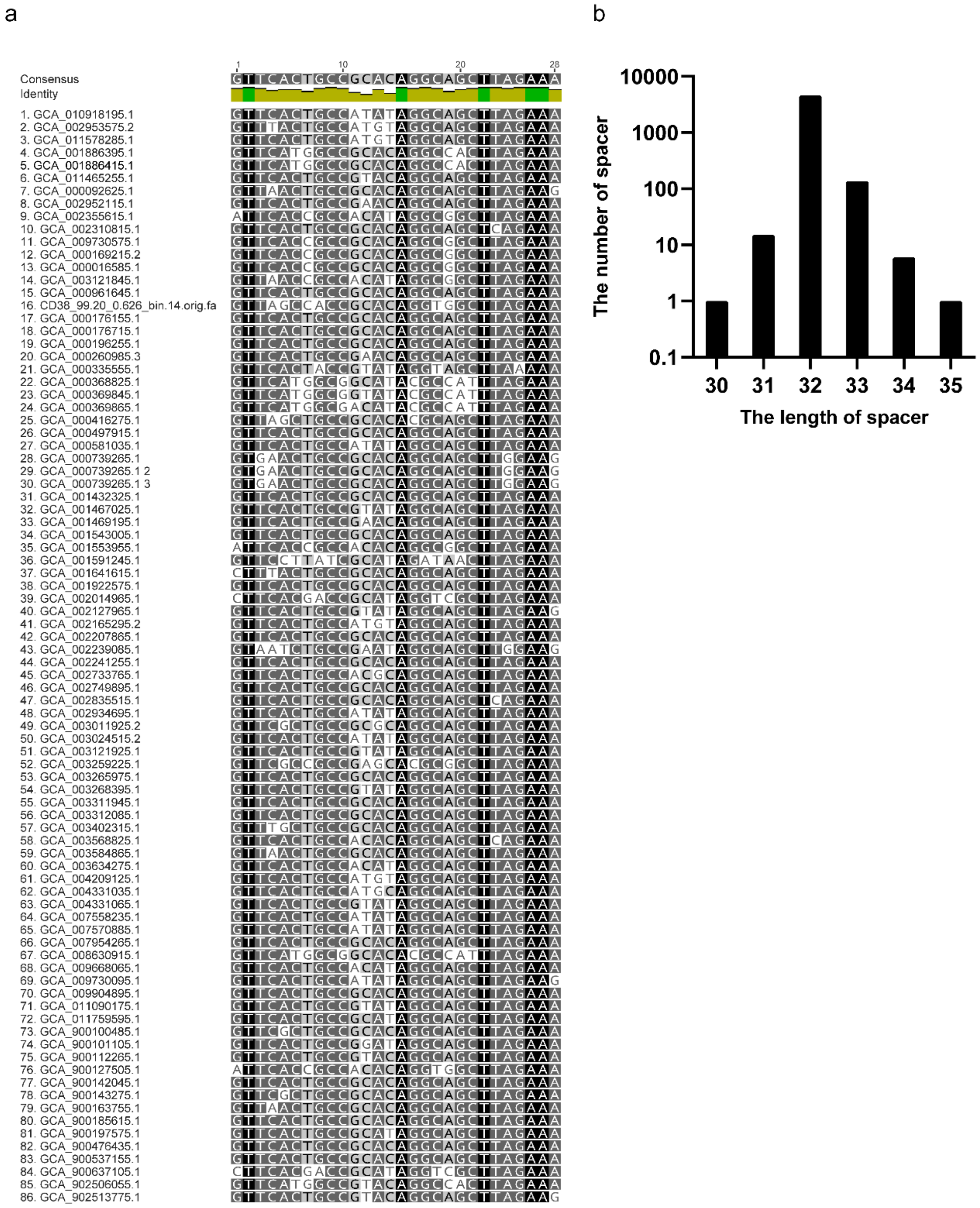


**Supplementary Fig. 2 | Analyses of CRISPR arrays next to the I-F2 Cas operon. a,** Multiple sequence alignment results of the repeats. CRISPR arrays were predicted in the vicinity of 86 Cas gene clusters from the 93 strains. The darker color at a position indicates a higher degree of conservation for the nucleotide. **b,** The total number of spacers of different lengths from CRISPR arrays next to the I-F2 Cas operons was counted and plotted in the bar graph.

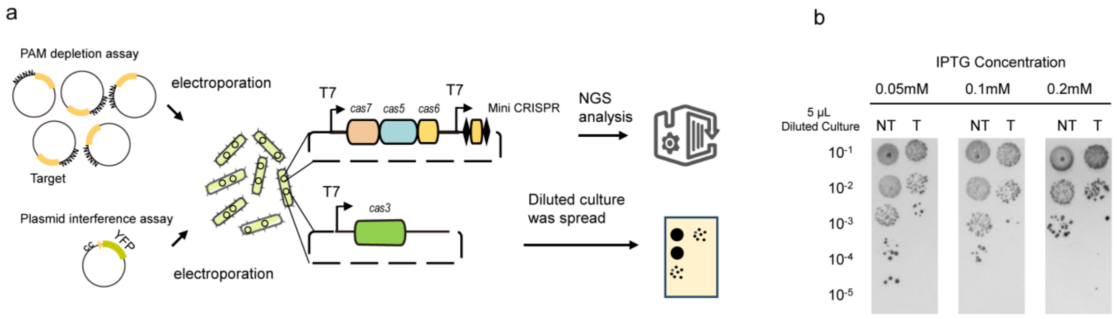


**Supplementary Fig. 3 | Mos350 I-F2 system provides immunity against plasmids containing protospacers with CC PAM at the 5′ end. a,** Schematic flow diagram of the PAM depletion assay and plasmid interference assay. **b,** Bacterial survival assay after induction of the Mos350 CRISPR-Cas system expression by different IPTG concentrations. NT: Plasmids expressing the Cascade-Cas3 without the mini-CRISPR; T: Plasmids expressing the Cascade-Cas3 and mini-CRISPR. The dilution gradient is shown on the left.

| Species | | | Abbreviation | Isolation source | The gene size of I-F2 Cascade (bp) |
|---|---|---|---|---|---|
| *Shewanella putrefaciens* CN32 | | ■ | SpuCN32 | Soil | 2511 |
| *Pseudoalteromonas tunicata* D2 | ■ | ■ | Ptu | Marine Broth | 2511 |
| *Alteromonas mediterranea* CP49 | | ■ | AmeCP49 | Seawater | 2565 |
| *Legionella longbeachae* B1445CHC | ■ | ■ | Llo | Homo sapiens | 2520 |
| *Methylophaga nitratireducenticrescens* JAM1 | ■ | ■ | MniJAM1 | Marine water | 2517 |
| *Moraxella osloensis* KMC41 | ■ | ■ | Mos41 | Cotton | 2574 |
| *Acinetobacter* sp. C16S1 | | ■ | AspC16S1 | Saline soil | 2601 |
| *Acinetobacter sp.* WCHAc010052 | ■ | ■ | AspWC | Sewage | 2559 |
| *Acinetobacter indicus* SGAir0564 | ■ | ■ | AinSGAir | Air | 2607 |
| *Acinetobacter* sp. 185 | ■ | ■ | Asp185 | Equus kiang | 2607 |
| *Moraxella osloensis* CCUG 350 | ■ | ■ | Mos350 | Human cerebrospinal fluid | 2709 |

■ Experiments in *E.coli*
■ Experiments in HEK293T

**Supplementary Fig. 4 | Source information of the I-F2 CRISPR-Cas systems utilized in the experiments.** The abbreviation of the strain name facilitates the description of the experimental results. Strain isolation sources were obtained from the BioSample database of NCBI. The eight systems expressed in *E. coli* are marked by orange squares and the 11 systems expressed in human cells are marked by blue squares. The gene size of Cascades was calculated as the total gene size of *cas5*, *cas6*, and *cas7*.

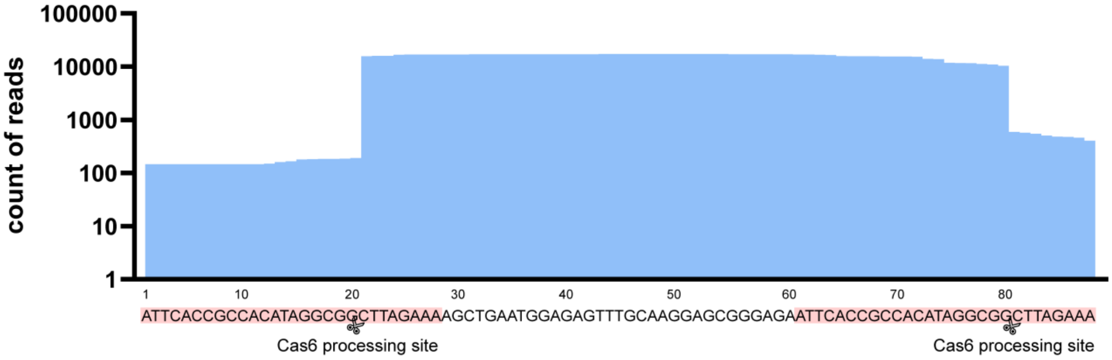


**Supplementary Fig. 5 | Small RNA sequencing reveals Cas6 cleavage site in repeat sequence.** The small RNA sequencing reads were mapped to the mini-CRISPR sequence with a spacer flanked by two repeats. The pink region represents the repeat sequence, and the Cas6 processing sites are marked by the scissors between positions 20 and 21 of the repeat.

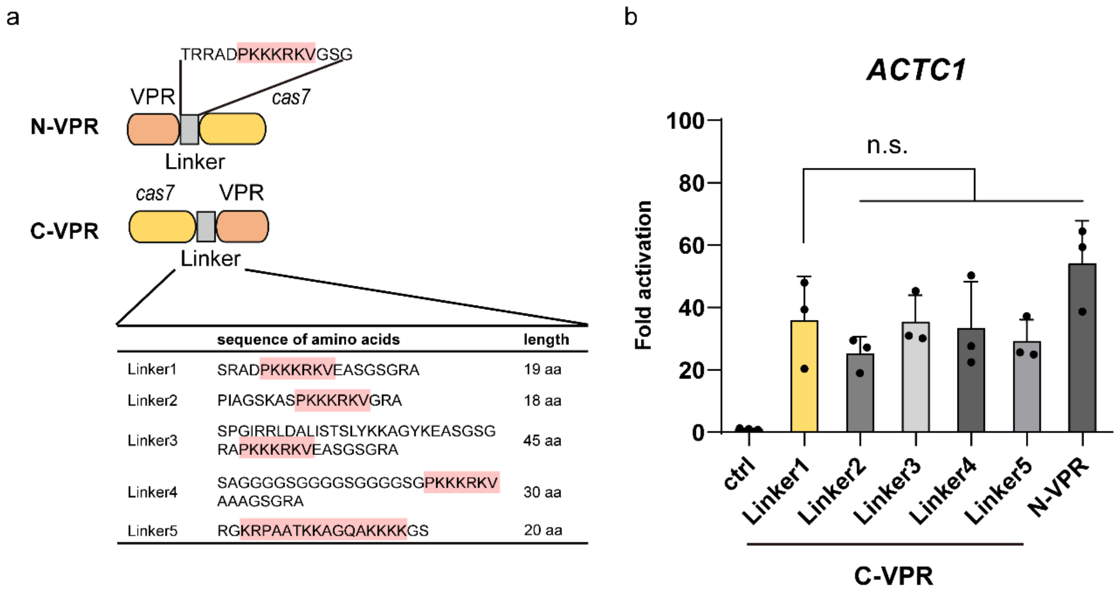


| | sequence of amino acids | length |
|---|---|---|
| Linker1 | SRADPKKKRKVEASGSGRA | 19 aa |
| Linker2 | PIAGSKASPKKKRKVGRA | 18 aa |
| Linker3 | SPGIRRLDALISTSLYKKAGYKEASGSGRAPKKKRKVEASGSGRA | 45 aa |
| Linker4 | SAGGGGSGGGGSGGGGSGPKKKRKVAAAGSGRA | 30 aa |
| Linker5 | RGKRPAATKKAGQAKKKKGS | 20 aa |

**Supplementary Fig. 6 | Optimizing transcriptional activation efficiency of Cascade-VPR with different linkers and gene arrangements. a,** Schematic representation of six designs for Mos350 Cas7-VPR fusion. The table shows the amino acid sequence information and length of different linkers, of which linker1 was the initial design. The pink color denotes the NLS. N-VPR: The VPR fused to the N-terminus of Cas7; C-VPR: The VPR fused to the C-terminus of Cas7. **b,** Comparison of six designs for transcriptional activation of the *ACTC1* gene in HEK293T cells. All values are represented as mean ± s.d. of three biological replicates. Statistical significance was assessed by one-way ANOVA (ns, not significant; $*P < 0.05$; $**P < 0.01$; $***P < 0.001$).

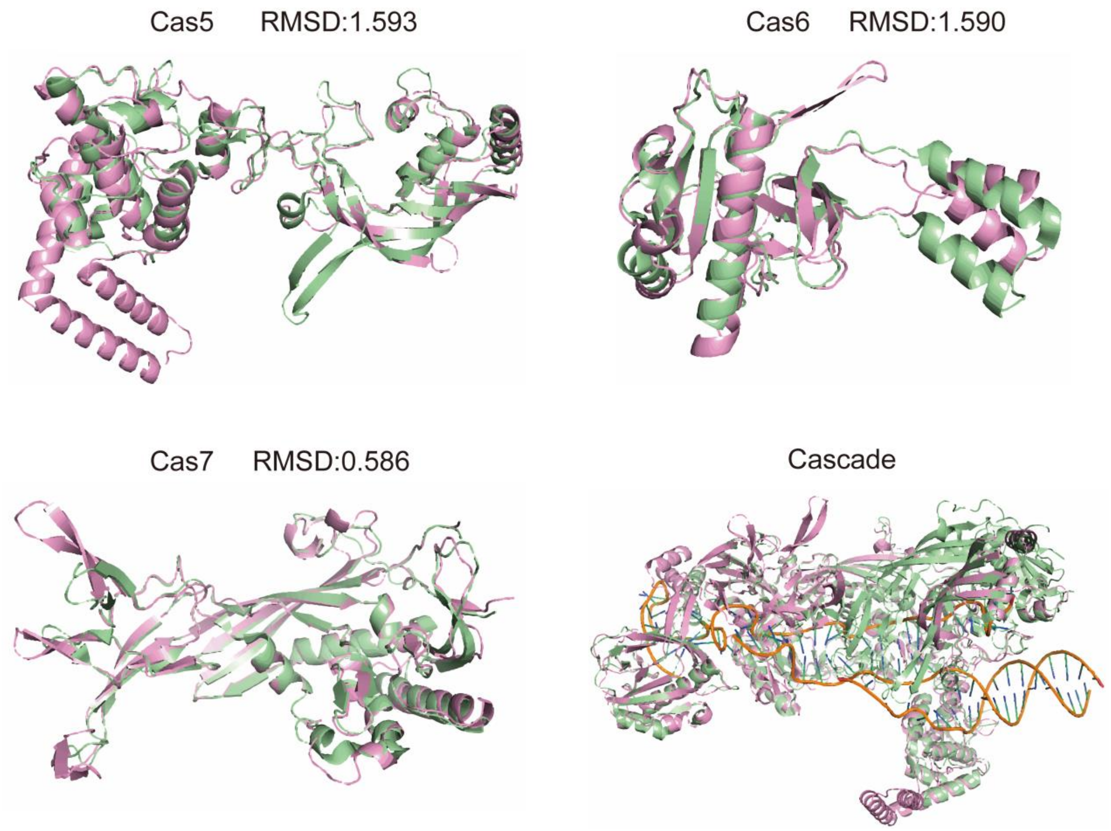


**Supplementary Fig. 7 | Superposition of three-dimensional structures of Cas5, Cas6, Cas7, and Cascade from Spu and Mos350 I-F2 systems.** The structures of the Spu Cascade and the individual subunits were derived from PBD 5O6U and rendered in pale green. The structures of all subunits of Mos350 were downloaded from AlphaFold and rendered pink. The subunit structures from two strains were aligned using Pymol, and the RMSD values were calculated.

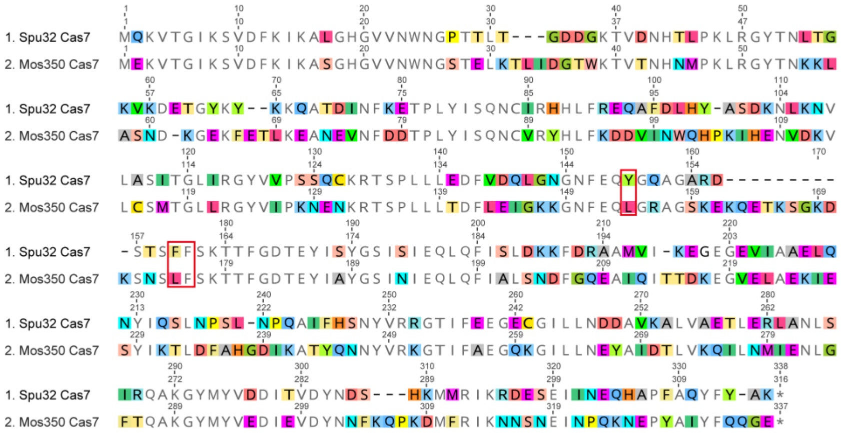


**Supplementary Fig. 8 | Sequence alignment between Spu Cas7 and Mos350 Cas7.** The red boxes indicate the aromatic residues emanating from the Cas7 thumb (Y149, F160, and F161), forming a stacking force in Spu I-F2, while Mos350 Cas7 possesses only one aromatic residue in the corresponding positions.

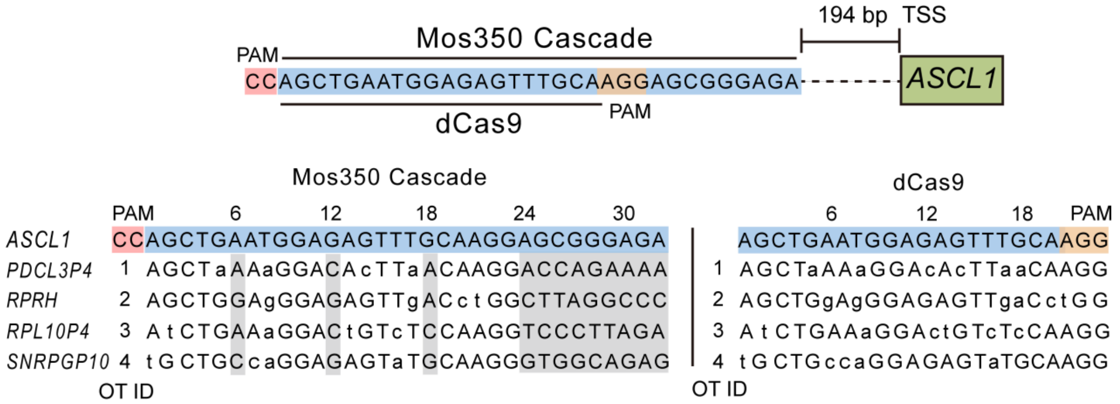


**Supplementary Fig. 9 | Predicted off-target sites of the Mos350 Cascade-VPR and dCas9-VPR targeting the *ASCL1* locus.** The blue regions mark the target sequences. The pink regions mark the PAM motifs of Cascade-VPR and the orange regions mark the PAM motifs of dCas9-VPR. Potential off-target sites and off-target gene names were listed. The gray area indicates that complementary pairing was not required. Lowercase indicates mismatched bases. OT ID: Off-target site ID.

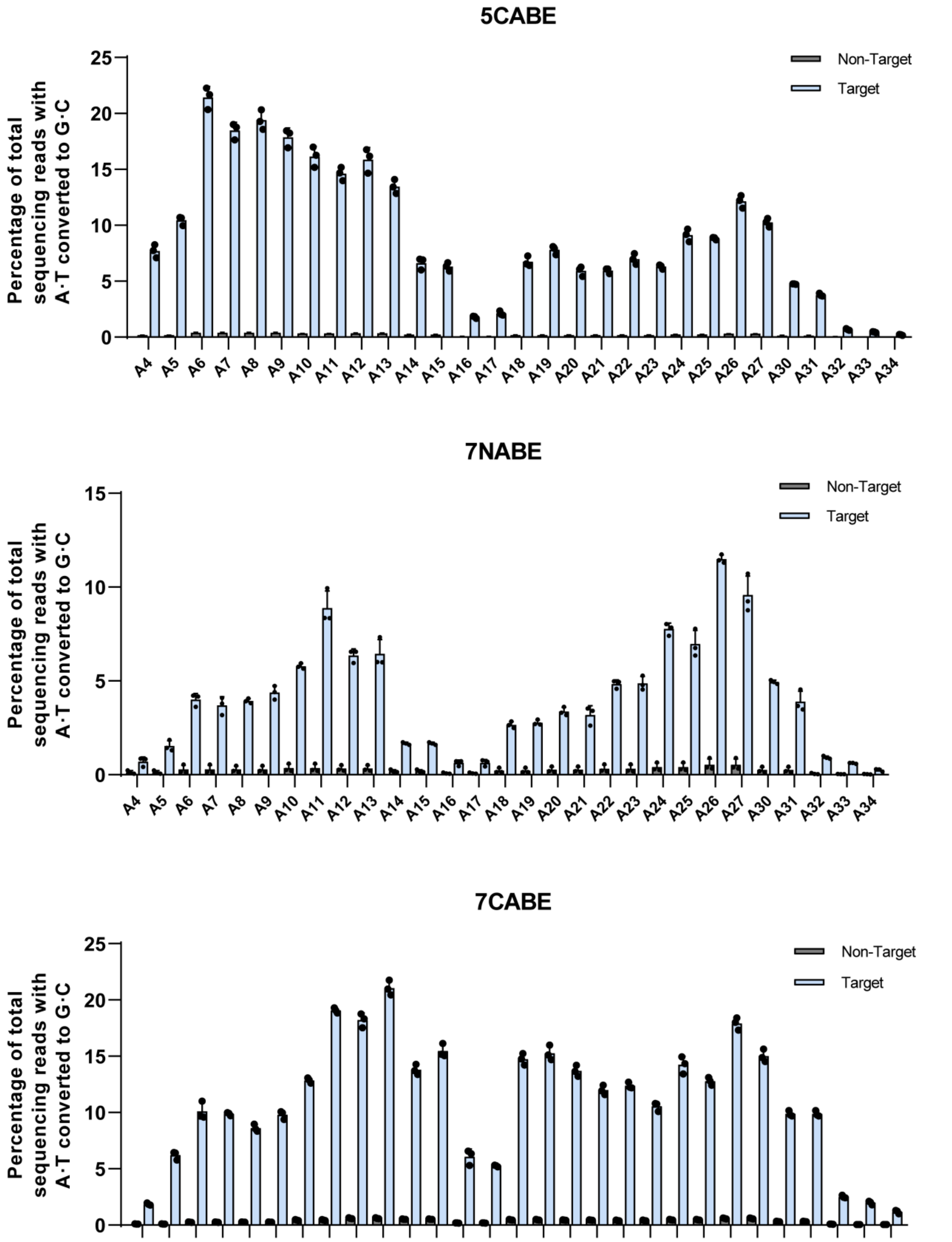


**Supplementary Fig. 10 | Base editing windows of three different I-F2 ABE constructions in human cells.** A·T-to-G·C editing windows of 5CABE, 7NABE, and 7CABE in HEK293T cells when aiming at AC-enriched targets on the *NIBAN* gene. The site 1 and site 2 on the *NIBAN* gene were enriched with AC bases. Non-Target: Only the plasmid expressing the Cascade-ABE was transfected. Target: The plasmid expressing the Cascade-ABE and the plasmid expressing the mini-CRISPR were co-transfected.

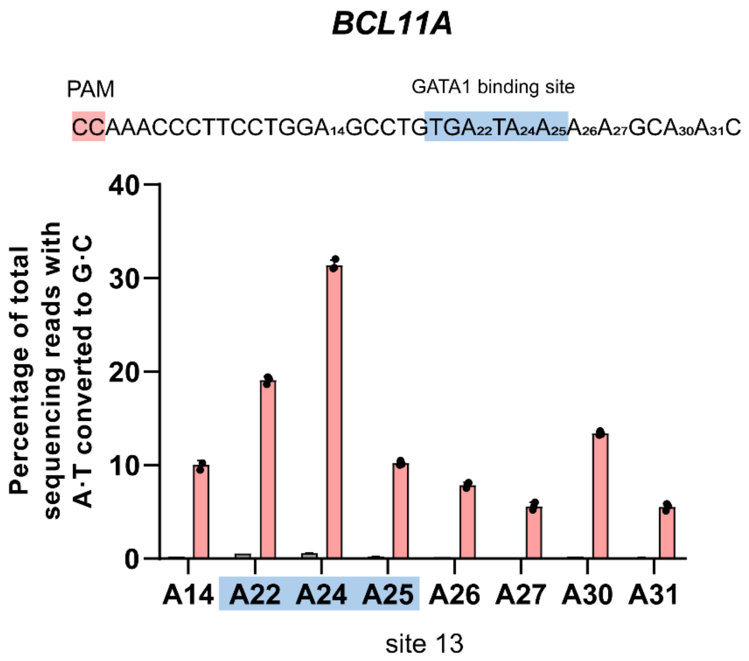


**Supplementary Fig. 11 | Base editing efficiency of 5NABE at the GATA1 site of the *BCL11A* enhancer.** The pink area marks the PAM of site 13, and the blue area labels the GATA1 binding site.

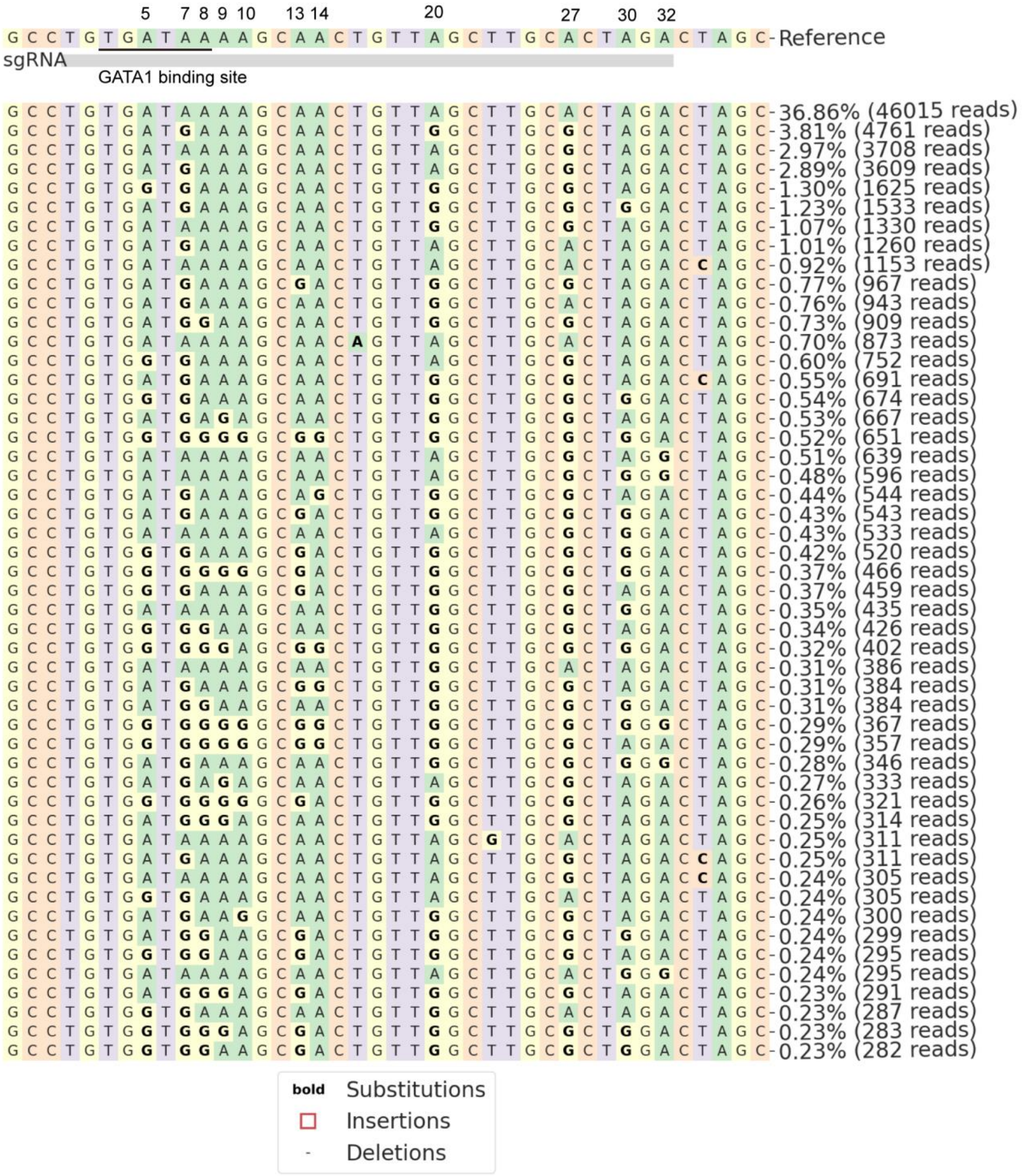


**Supplementary Fig. 12 | Allele compositions following treatment with 5NABE at the GATA1 binding site of the *BCL11A* enhancer.** Since 5NABE is a wide-window editing tool, in addition to containing A-T-to-G-C edited alleles at positions 5, 7, and 8 (site 12), the enhancer region excluding the GATA1 binding site also produced substantial A·T-to-G·C editing, which might further contribute to the suppression of *BCL11A* expression.

a

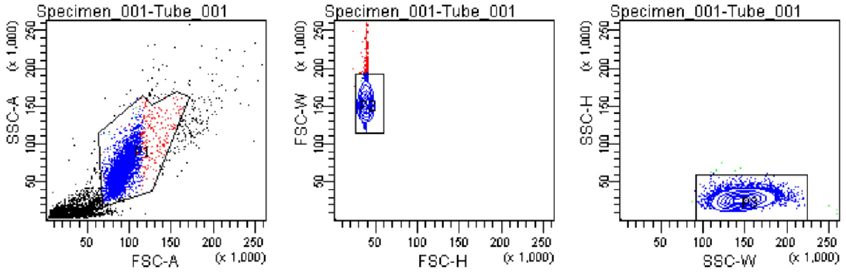


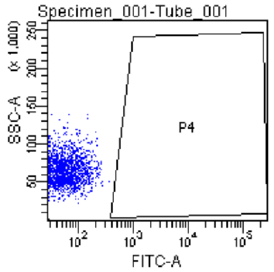


Tube: Tube_001

| Population | #Events | %Parent | %Total |
|---|---|---|---|
| All Events | 10,000 | #### | 100.0 |
| P1 | 6,599 | 66.0 | 66.0 |
| P2 | 6,358 | 96.3 | 63.6 |
| P3 | 6,348 | 99.8 | 63.5 |
| P4 | 0 | 0.0 | 0.0 |

b

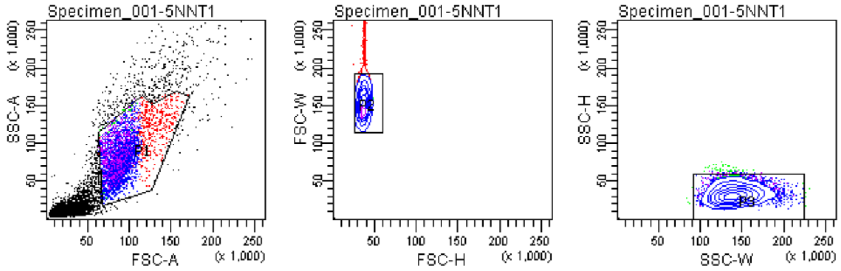


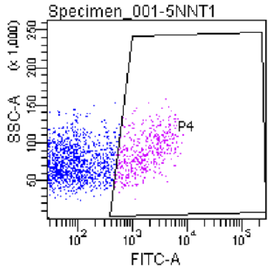


Tube: 5NNT1

| Population | #Events | %Parent | %Total |
|---|---|---|---|
| All Events | 10,000 | #### | 100.0 |
| P1 | 4,271 | 42.7 | 42.7 |
| P2 | 3,777 | 88.4 | 37.8 |
| P3 | 3,729 | 98.7 | 37.3 |
| P4 | 495 | 13.3 | 5.0 |

c

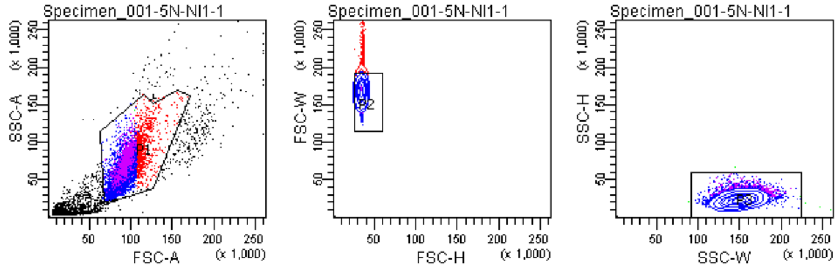


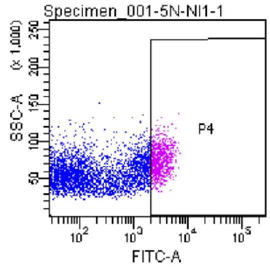


Tube: 5N-NI1-1

| Population | #Events | %Parent | %Total |
|---|---|---|---|
| All Events | 10,000 | #### | 100.0 |
| P1 | 7,487 | 74.9 | 74.9 |
| P2 | 6,469 | 86.4 | 64.7 |
| P3 | 6,463 | 99.9 | 64.6 |
| P4 | 798 | 12.3 | 8.0 |

**Supplementary Fig. 13 | FACS gating examples for EGFP-positive cells sorting conditions. a,** EGFP-negative cells were never collected under the sorting conditions. FITC ≥ 500. **b,** EGFP-positive cells were collected for base editing efficiency analysis. FITC ≥ 500. **c,** EGFP-positive cells with higher expression were collected for base editing efficiency analysis. FITC ≥ 2000.

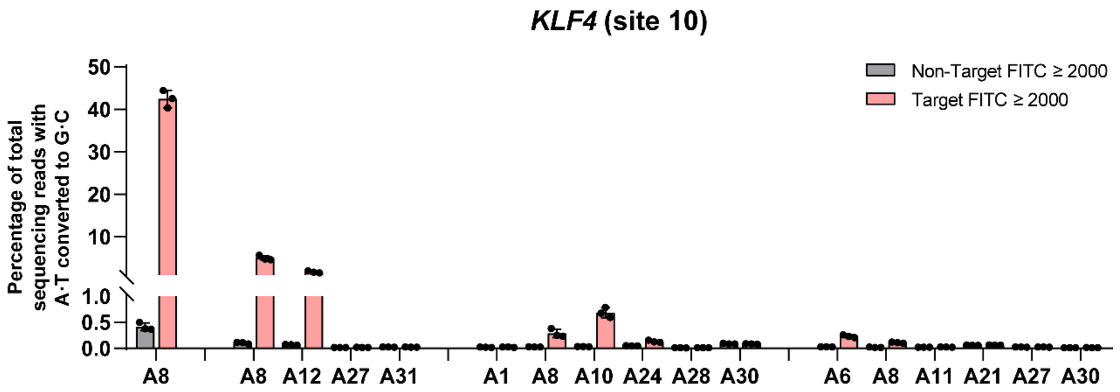


**Supplementary Fig 14 | Off-target efficiencies of 5NABE at site 10 with a higher fluorescence threshold.** Base editing efficiency of 5NABE at target (Fig. 6d) and off-target sites when cells with FITC ≥ 2000 were collected.
