## Supplementary material for "Engineered minimal type I CRISPR-Cas system for transcriptional activation and base editing in human cells": Custom scripts

### Supplementary Note

**Contents:**

1. The script used to calculate the coverage of sRNA-seq reads from BLAST output.
2. The script used to split fastq barcode samples.

```
#!/usr/bin/perl -w
#This script is used to calculate the coverage of sRNA-seq reads from BLAST output.

use strict;
use Getopt::Long;

my %opts;
GetOptions(\%opts,"i=s","b=s","l=i","o=s","h");
if (!(defined $opts{i} and defined $opts{b} and defined $opts{o}) || defined $opts{h}) {
		#necessary arguments
	&usage;
}

my $R2reads_blastout_file=$opts{'i'};
my $R1reads_blastout_file=$opts{'b'};
my $fileout=$opts{'o'};



my @array=();
my $targetlen;
if(defined($opts{'l'})){
	$targetlen=$opts{'l'};
}
else{
	$targetlen=400;
}

@array = ( split //, 0 x $targetlen );



open IN,"<$R2reads_blastout_file" or die $!;
open OUT,">$fileout";
# E116	74	1	74	174	247	756	147	2e-40	74/74	100	pTA-Hubei-tRNA		R1
# E142	123	1	123	174	296	756	244	2e-69	123/123 100	pTA-Hubei-tRNA		R1
# E166	132	1	132	174	305	756	262	1e-74	132/132 100	pTA-Hubei-tRNA		R1
# E201	49	2	49	528	575	756	95.6	5e-25	48/48	100	pTA-Hubei-tRNA		R1
# E246	110	1	110	178	287	756	218	1e-61	110/110 100	pTA-Hubei-tRNA
my (@ab,%r2,%startr2,%endr2);
while(<IN>){
	chomp;
	@ab = split(/\s+/,$_);
	$r2{$ab[0]} = $_;
```

```
	$startr2{$ab[0]} = $ab[4];
	$endr2{$ab[0]} = $ab[5];

}


open IN,"$R1reads_blastout_file" or die $!;
my (@a);
while(<IN>){
	chomp;
	@a = split(/\s+/,$_);
	if (exists($r2{$a[0]})){
		print "r1 have hit in r2:\n$_\n$r2{$a[0]}\n";
		my @temp = sort {$a<=>$b} ($a[4],$a[5],$startr2{$a[0]},$endr2{$a[0]});
		my $min = $temp[0];
		my $max = $temp[-1];
		print "before:$a[4],$a[5],$startr2{$a[0]},$endr2{$a[0]}\n";
		print "pair-end map sort:", join(",,",@temp),",r1 r2:min\t$min\tmax\t$max\n";

		for (my $i=$min;$i<=$max;$i++){
			$array[$i]++;
		}
	}
	else{
		print "r1 no hit in r2:$_,,single map:start\t$a[4]\tend\t$a[5]\n";
		my @temp = sort {$a<=>$b} ($a[4],$a[5]);
		my $min = $temp[0];
		my $max = $temp[-1];
		print "single map sort:", join(",,",@temp),",r1:min\t$min\tmax\t$max\n";
		for (my $i=$min;$i<=$max;$i++){
			$array[$i]++;
		}
	}

}



for (my $j=1;$j<$targetlen;$j++){
	print OUT "$j\t$array[$j]\n";

}
```

```
sub usage{
	print <<"USAGE";

Usage:
	$0 -i <input file> -b <input file> -l <input length> -o <output file>
	perl	sRNA-seq-coverage.pl	-i	MOS350-7v-AS1.trimmed.R22target.bln.out	-b
MOS350-7v-AS1.trimmed.R12target.bln.out	-l 200 -o	MOS350.cov > MOS350.cov.detail

options:
	-i input R2reads_blastout_file:
	for example:
# Query name	Letter	QueryX	QueryY	SbjctX	SbjctY	Length	Score	E value Overlap/total	Identity	Sbject
	E713	59	3	59	77	21	88	106	4e-29	57/57	100	DRspacerDR
	E3882	60	1	60	80	21	88	111	9e-31	60/60	100	DRspacerDR
	E4090	59	3	59	77	21	88	106	4e-29	57/57	100	DRspacerDR
	E4384	60	1	60	80	21	88	111	9e-31	60/60	100	DRspacerDR

	-b input R1reads_blastout_file:
	for example:
# Query name	Letter	QueryX	QueryY	SbjctX	SbjctY	Length	Score	E value Overlap/total	Identity	Sbject
	E713	59	1	57	21	77	88	106	4e-29	57/57	100	DRspacerDR
	E3882	60	1	60	21	80	88	111	9e-31	60/60	100	DRspacerDR
	E4090	59	1	57	21	77	88	106	4e-29	57/57	100	DRspacerDR
	E4384	60	1	60	21	80	88	111	9e-31	60/60	100	DRspacerDR

	-l target length :Integer default is 400nt
	-o output file

	-h help
USAGE
	exit(1);
}
```

```
#!/usr/bin/perl -w
#This script is used to split fastq barcode samples.

#use strict;
use Getopt::Long;

my %opts;
GetOptions(\%opts,"i=s","b=s","h");
if (!(defined $opts{i} and defined $opts{b}) || defined $opts{h}) {			#necessary
arguments
	&usage;
}

my $barcodefile=$opts{'i'};
my $flash_merge_file=$opts{'b'};



open IN, "$barcodefile" or die $!;
my (%barcodegroup2sample,%groupsample);

while (<IN>){
	chomp;
	my @a = split(/\t/,$_);
	$barcodegroup2sample{$a[2]}{$a[0]}=$a[1];
	$groupsample{$a[0]}{$a[1]}=1;
	#print "f1:$a[0],,$a[1]\n";
}


open IN, "$flash_merge_file" or die $!;


my $groupname=$flash_merge_file;
$groupname=~s/(\S+).extendedFrags.fastq/$1/;


foreach my $s (keys %{$groupsample{$groupname}}){
	my $oname = $groupname."-".$s;
	print "outfilename:$oname\n";
	open $oname,">$groupname"."-"."$s".".fq" or die $!;
}

my $reads_seq;
```

```
my ($out,$flag,$seq);
my $aline;
my $plus_line;
my $reads_quality;
while( $aline=<IN>){
	$reads_seq=<IN>;
	$flag =0;
	$seq = "";
	foreach my $k (keys %barcodegroup2sample){

		if ($reads_seq =~/^$k/) {
			#print "$k,,$reads_seq\n";
			$flag =1;
			$seq = $k;
			last;
		}
	}
	if ($flag){
		$out = $groupname."-".$barcodegroup2sample{$seq}{$groupname};
		#print "exist,$seq,,$hash{$seq},,$out,,$reads_seq,,\n";
		print $out "$aline";
		my $rmbarseq = substr($reads_seq,6);
		print $out "$rmbarseq";
		#print $out "$reads_seq";
		$plus_line=<IN>;
		print $out "$plus_line";
		$reads_quality=<IN>;
		my $rmbarseqqual = substr($reads_quality,6);
		print $out "$rmbarseqqual";
	}else{
		<IN>;
		<IN>;
	}

}
close IN;



sub usage{
	print <<"USAGE";

Usage:
	$0 -i <input file> -b <input file>
```

```
	perl  fastq-sample-split-multipleGroups.pl  -i  group-sample-barcode.txt  -b C1.extendedFrags.fastq
options:
	-i input barcodefile
	#example:sequce file name	sample name barcode
	# C1	188-1	CTTGTA  GAGGAGTGTTCAGTCTCCGTGAACGTTCCCTTAGCAC
	AGCTGCTCACTTGAGCCTCTGGGTCTAGAACCCTCTGGGGACCGTTTGAGGAGTGTTCAGT
CTCCGTGAACGTTCCCTTAGCACTCTGCCACTTATTGGGTCAGCTGTTAACATCAGTACGTTAAT
GTTTCCTGATGGTCCATGTCTGTTACTCGCCTGTCAAGTGGCGTGACACCGGGCGTGTTCCCCA
GAGTGACT
	# C1	191-1	TGACCA GGGAGGTCAGAAATAGGGGGTCCAGGAGCAAACTCCC
	TCTCTGTACATGAAGCAACTCCAGTCCCAAATATGTAGCTGTTTGGGAGGTCAGAAATAGG
GGGTCCAGGAGCAAACTCCCCCCACCCCCTTTCCAAAGCCCATTCCCTCTTTAGCCAGAGCCGG
GGTGTGCAGACGGCAGTCACTAGGGGGCGCTCGGCCACCACAGGGAAGCTGGGTGAATGGAG
CGAGCA

	-b input flash_merge_file: this file is the output of FLASH(Fast Length Adjustment of SHort reads) software, which  is an accurate and fast tool to merge paired-end reads that were generated from DNA fragments whose lengths are shorter than twice the length of reads.

	-h help
USAGE
	exit(1);
}
```
